# DHA deficiency drives neural oxidized-lipid stress and microglial activation

**DOI:** 10.64898/2026.09.22.753588

**Authors:** Ryo Hagimori, Purnima Gogoi, Pawan K. Shahi, Ryoya Ueno, Greg Barrett-Wilt, Michael Landowski, Kazuya Oikawa, Madison Tytanic, Richard S. Brush, Jingyang Wei, Maya S. Guerriero, Gillian J. McLellan, Sakae Ikeda, Martin-Paul Agbaga, Bikash R. Pattnaik, Ken-ichi Yamada, Akihiro Ikeda

## Abstract

Despite the well-established importance of DHA in neural health, the mechanisms by which neural tissues respond to declining DHA availability and how this response contributes to aging remain poorly understood. Here, we show that chronic DHA deficiency triggers a neural-specific compensatory lipid-remodeling program with pathological consequences. DHA loss causes a neural-tissue-selective shift from DHA-containing phospholipids toward arachidonic and adrenic acid-containing ω-6 species, accompanied by an RPE-associated polyunsaturated fatty acid biosynthetic program. This remodeling expands the pool of oxidation-prone lipids, redirects lipid peroxidation toward ω-6-derived products, and increases oxidative damage. Dietary DHA restoration reversed lipid remodeling, oxidative stress, inflammation, and visual dysfunction, establishing DHA deficiency as a causal driver of these phenotypes in mice. We identified an APOE-dependent oxidized-lipid disposal pathway that transfers oxidized lipid cargo from the neural retina to subretinal microglia, limiting its retention within the neural retina and protecting against degeneration. However, with persistent lipid uptake, oxidized phospholipids accumulate in microglial lysosomes, with sustained galectin-3 activation. Galectin-3 deficiency preferentially protected against later-stage degeneration, indicating that prolonged lipid burden converts the initial clearance response into a pathogenic response. Physiologically aged retinas also showed a similar shift toward ω-6 lipid accumulation. These findings define an adaptive-to-maladaptive lipid-remodeling–microglia axis linking declining DHA availability to age-related neural dysfunction.

## Introduction

Aging profoundly alters lipid metabolism and innate immune function^1,2,3^, yet how these processes interact to drive functional decline in neural tissues during aging remains poorly understood. The retina is especially vulnerable to such age-related changes because photoreceptors contain exceptionally lipid-rich membranes that undergo continuous renewal and are exposed to sustained metabolic and oxidative stress^4^. Docosahexaenoic acid (DHA; 22:6, ω-3) is highly enriched in neuronal membrane phospholipids, particularly those of photoreceptor outer segments, where it supports the membrane properties required for visual function^4,5^. Retinal DHA availability declines with aging^6^, but whether such loss actively remodels neural membrane lipids and contributes to age-related dysfunction remains unknown.

Lipid peroxidation provides a potential link between altered membrane composition and neural tissue damage^7,8^. Reactive oxygen species can oxidize polyunsaturated fatty acids (PUFAs) within membrane phospholipids, generating peroxidized lipid species and secondary reactive products that modify cellular macromolecules and engage inflammatory pathways^9,10^. The retina is particularly exposed to such reactions because of its high oxygen demand, continuous light exposure, and abundance of PUFA-containing membranes^11^, making it particularly well suited for examining how age-related changes in membrane composition influence oxidative lipid stress. Crucially, membrane fatty-acid composition is a key determinant of which oxidized lipids are produced. A decline in DHA may therefore have consequences beyond the loss of a structural membrane component: compensatory replacement of DHA-containing phospholipids by other PUFA-containing species could reshape the substrates available for peroxidation and thereby alter the identity and distribution of oxidized lipids generated during stress. However, whether declining DHA availability triggers such remodeling within neural tissue, where it is spatially organized and how it reshapes oxidative lipid stress remain unknown.

Cellular responses to lipid peroxidation extend beyond the cells in which oxidative damage originates. Emerging evidence indicates that stressed neurons can export excess or peroxidized lipids to neighbouring glial cells^12,13^, where their sequestration in lipid droplets and subsequent metabolism can limit neuronal lipid toxicity^13,14^. Such intercellular lipid transfer may therefore constitute an adaptive mechanism for maintaining neural lipid homeostasis. However, lipid handling by glial cells can also become maladaptive: peroxidized lipids released from damaged neurons can accumulate in microglia and engage inflammatory pathways that promote neurodegeneration^15^. In the retina, microglia undergo a characteristic redistribution from their normal plexiform-layer niches into the subretinal space during aging and degeneration^16,17,18^. This spatially defined response makes the retina particularly well suited for determining how lipid trafficking shapes microglial behavior during neural aging. This stereotyped redistribution of retinal microglia therefore offers a means to address these unresolved questions: whether their movement into the subretinal space is coupled to intercellular lipid trafficking, which pathways mediate lipid transfer, and whether such lipid handling protects or damages neural tissue.

Here, using *Tmem135* mutant mice, which exhibit chronic systemic DHA deficiency and accelerated retinal aging phenotypes^18,19,20^, we combined dietary DHA restoration with tissue-resolved lipidomics, oxidized-lipid profiling and genetic perturbation of lipid-responsive microglial pathways. We show that DHA deficiency does not simply deplete polyunsaturated lipids but instead drives a neural-tissue-selective replacement of DHA-containing phospholipids with ω-6-containing species, accompanied by a retinal pigment epithelium (RPE)-associated biosynthetic program. Importantly, dietary DHA supplementation restored the retinal lipid environment and rescued the associated age-related phenotypes, establishing DHA loss as a causal driver of this process. This altered lipid environment redirects lipid peroxidation towards ω-6-derived products, resulting in their preferential accumulation within the neural retina. We further identify an apolipoprotein E (APOE)-dependent pathway that transfers oxidized lipids from the neural retina to subretinal microglia and limits their retention within neural tissue. At later stages of retinal aging, however, sustained lipid-associated lysosomal stress engages a galectin-3-dependent microglial program that contributes to neurodegeneration. Finally, physiological aging recapitulates this lipid-remodeling signature, which is further amplified by DHA deficiency. Together, these findings establish declining DHA availability as a causal driver of a lipid-remodeling–oxidation–microglia axis that links neural membrane composition to age-related tissue dysfunction.

## Results

### DHA deficiency drives broad aging-related retinal pathology

DHA is highly enriched in the neural retina and supports photoreceptor membrane integrity and visual function. Retinal DHA availability declines with age^6^, and the extent to which this decline contributes to aging-related retinal dysfunction and neuroinflammation remains unclear. TMEM135, a peroxisome-localized transmembrane protein, has been implicated in the terminal handling of DHA synthesized through the Sprecher pathway, likely by facilitating the export of newly generated DHA from peroxisomes^19,21^. *Tmem135* mutant mice exhibit a marked reduction in DHA across multiple tissues (Supplementary Fig. 1)^19^ and develop a broad spectrum of age-related retinal abnormalities^18^, offering a unique opportunity to determine how chronic DHA deficiency influences retinal aging. We tested whether restoring DHA availability with a DHA-rich fish-oil diet could reverse the structural, functional and inflammatory retinal abnormalities in *Tmem135* mutant mice.

We fed *Tmem135* mutant mice either a control diet (CD) or a diet supplemented with DHA-rich fish oil (10% w/w, FOD). LC–MS/MS-based lipidomic profiling showed that the fish-oil diet broadly restored DHA-containing lipid species toward wild-type (WT) levels in *Tmem135* mutant retinas (Fig. 1a). Histological analyses demonstrated that dietary fish-oil substantially mitigated retinal degeneration in these mice (Fig. 1b–d). We then assessed whether these structural improvements were accompanied by recovery of retinal function using electroretinography. Under scotopic conditions, fish-oil supplementation restored the a-wave amplitude, reflecting rod photoreceptor activity, and the b-wave amplitude, representing rod-driven bipolar-cell responses with contributions from Müller glia toward wild-type levels, which were reduced in *Tmem135* mutant mice (Fig. 1e–h). Photopic recordings showed similar reductions and restoration of cone-mediated retinal responses (Supplementary Fig. 2a–d). The c-wave, which largely reflects the RPE cell activity in response to light-evoked changes in subretinal potassium concentrations, was also impaired in *Tmem135* mutant mice and substantially recovered in fish-oil-fed mutants (Fig. 1i). Microglial activation and the accumulation of myeloid cells in the subretinal space are prominent features of retinal aging and age-related retinal disease^16,18^. *Tmem135* mutant mice exhibited increased number of IBA1⁺ microglia within the inner plexiform layer (IPL) and outer plexiform layer (OPL)^18^ (Supplementary Fig. 2e), where retinal microglia normally reside, alongside the marked accumulation of IBA1⁺ myeloid cells in the subretinal space (Fig. 1j,k, Supplementary Fig. 2e). DHA-rich fish-oil supplementation largely suppressed this subretinal myeloid-cell accumulation (Fig. 1j,k), indicating that DHA restoration attenuates the age-related inflammatory phenotype of the *Tmem135* mutant retina. Together, these findings show that a DHA-rich fish oil diet improves photoreceptor functions, inner retinal circuits, and the RPE cell functions. To define the molecular pathways associated with photoreceptor degeneration under conditions of DHA depletion, we performed RNA sequencing (RNA-seq) of neural retinas from WT and *Tmem135* mutant mice maintained on either a control diet (CD) or a DHA-rich fish-oil diet (FOD). Gene set enrichment analysis (GSEA) revealed enrichment of multiple stress-associated pathways in *Tmem135* mutant retinas, including endothelin (EDN)-mediated stress signaling^22^, oxidative stress responses and complement activation (Fig.1 l,m,n), and FOD supplementation largely reversed these stress-related pathways (Fig. 1o, Supplementary Fig. 2f,g). Consistent with these transcriptional changes, *Tmem135* mutant retinas exhibited pronounced Müller glial activation (Supplementary Fig. 2h,i) and increased expression of the central complement component C3 in Müller cells (Fig. 1o, Supplementary Fig. 2j), both of which were attenuated by fish-oil supplementation (Fig. 1o). Moreover, activated Müller glia extended processes that closely contacted microglia in the IPL/OPL under DHA depletion (Supplementary Fig. 2k), suggesting that Müller glia may relay neural retinal stress to microglia^23,24^, thereby contributing to their activation and subsequent migration towards the subretinal space. Consistent with this possibility, microglial activation was evident within the IPL and OPL, where retinal microglia normally reside, and was accompanied by a prominent accumulation of IBA1⁺ myeloid cells in the subretinal space (Supplementary Fig. 2e). To determine whether these subretinal cells originated from resident retinal microglia, we fate-mapped *Tmem119*-expressing microglia using the *Tmem119^CreERT2^*;*Rosa26^mTmG^* reporter strain, in which tamoxifen-induced recombination permanently switches membrane fluorescence from tdTomato to GFP^25^ (Supplementary Fig. 2l,m). GFP⁺ cells accumulated progressively in the subretinal space with age compared to the time point immediately after tamoxifen administration, demonstrating that resident retinal microglia migrate into this compartment in *Tmem135* mutant mice (Supplementary Fig. 2n).

**Figure 1.**
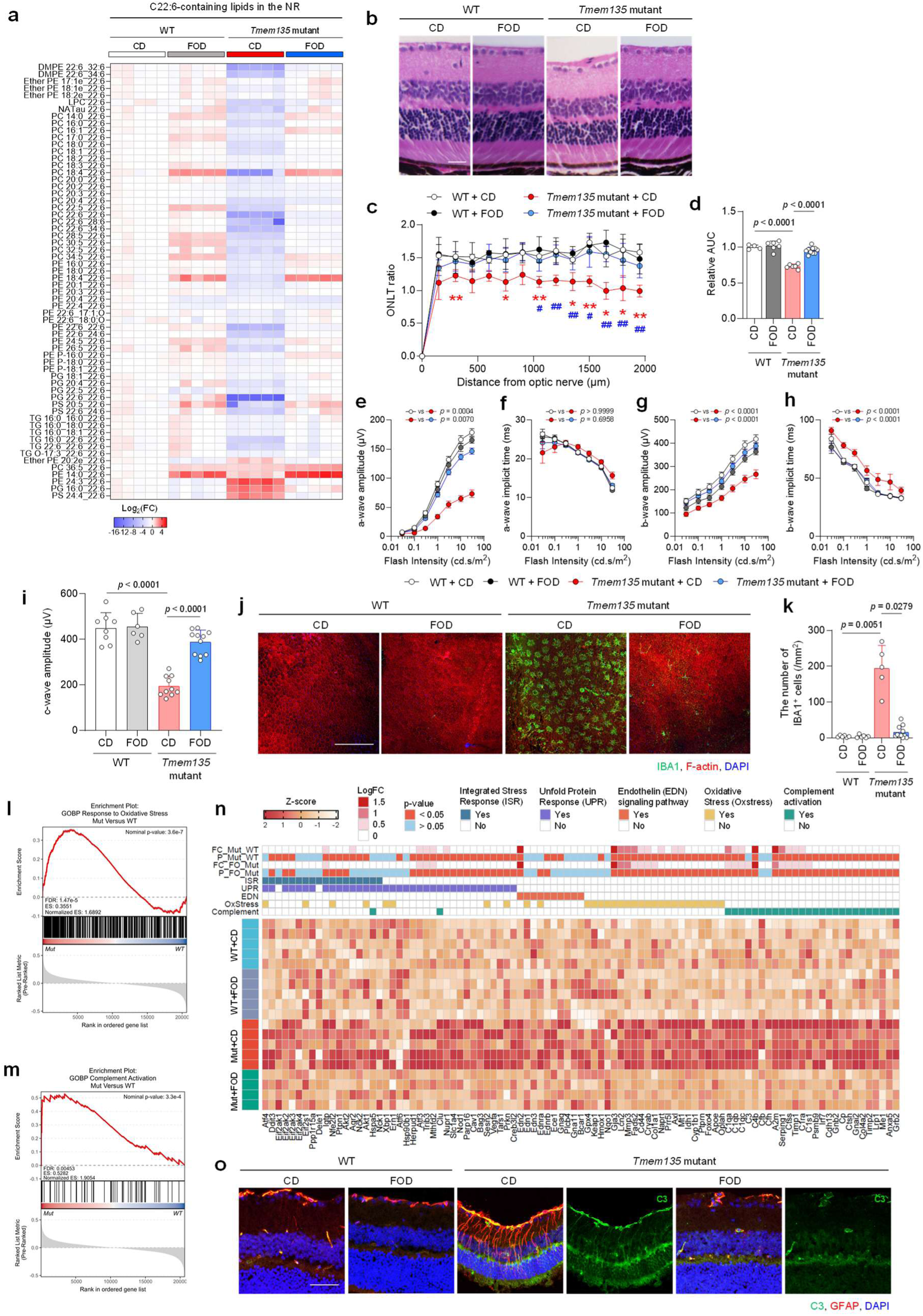
DHA deficiency drives broad aging-related retinal pathology. **a**, Heatmap of DHA (22:6)-containing lipids in the neural retina of WT and *Tmem135* mutant mice fed a control diet (CD) or fish-oil diet (FOD). **b**, Representative hematoxylin and eosin (H&E) staining of retinal sections. Scale bar, 25 μm. **c**, Outer nuclear layer thickness (ONLT) ratios. **d**, Area under the curve (AUC) of the ONLT ratio profiles. **e–i**, Quantification of scotopic ERG responses, showing a-wave amplitude (**e**), a-wave implicit time (**f**), b-wave amplitude (**g**), b-wave implicit time (**h**), and c-wave amplitude (**i**). **j**, Representative eyecup flat-mount images showing IBA1⁺ myeloid cells (green), F-actin (red) and DAPI-stained nuclei (blue) in the subretinal space. Scale bar, 200 μm. **k**, Quantification of IBA1⁺ myeloid cell numbers in the subretinal space. **l**, GSEA enrichment plot for the response to oxidative stress gene set in *Tmem135* mutant versus WT neural retina. **m**, GSEA enrichment plot for the complement activation gene set in *Tmem135* mutant versus WT neural retina. **n**, Heatmap showing expression of genes involved in the integrated stress response (ISR), unfolded protein response (UPR), endothelin (EDN)-related signaling, oxidative stress and complement activation. **o**, Representative retinal sections showing GFAP (red) and C3 (green) immunoreactivity. Scale bar, 50 μm. Data in (**c**,**d**,**k**) are presented as the mean ± s.d., whereas data in (**e–i**) are presented as the mean ± s.e.m. For ERG analyses (**e–i**), responses from both eyes were averaged to obtain one value per mouse. In the bar graphs, each data point represents an individual mouse. Sample sizes were n = 4–10 mice in (**c**,**d**), n = 6–11 mice in (**e–i**) and n = 5–11 mice in (**k**). For lipidomic (**a**) and RNA-seq (**l–n**) analyses, n = 4–5 mice per group. Statistical significance was assessed using two-way ANOVA followed by Tukey’s multiple-comparisons test (**c**) or Šídák’s multiple-comparisons test (**d**,**i**,**k**). For (**e–h**), statistical significance was assessed using two-way ANOVA with repeated measures on flash intensity and the Geisser–Greenhouse correction, followed by Šídák’s multiple-comparisons test of group marginal means. In (**c**), red asterisks indicate comparisons between WT + CD and *Tmem135* mutant + CD, whereas blue signs indicate comparisons between *Tmem135* mutant + CD and *Tmem135* mutant + FOD. One and two symbols denote *P* < 0.05 and *P* < 0.01, respectively. Exact *P* values are shown in the corresponding panels; values below 0.0001 are reported as *P* < 0.0001.

Together, these findings support a model in which DHA deficiency initiates photoreceptor and oxidative stress, leading to Müller glial and complement activation and the subsequent activation and migration of retinal microglia. Restoration of DHA availability suppresses the stress–glial response and preserves retinal structure and function.

### DHA deficiency induces local AA/AdA phospholipid accumulation in the neural retina

Because DHA is highly enriched in neural retinal membranes, particularly in photoreceptor outer segments^4,5,11^, we next asked how chronic DHA deficiency alters the phospholipid composition of the neural retina. Lipidomic profiling revealed broad increases in phospholipids containing the ω-6 polyunsaturated fatty acids arachidonic acid (AA; 20:4) and adrenic acid (AdA; 22:4) across multiple lipid subclasses in *Tmem135* mutant mice (Fig. 2a,b and Supplementary Figs. 3a and 4a–c). A similar enrichment of AA- and AdA-containing phospholipids was observed in the brain, indicating that this response is shared across neural tissues rather than restricted to the retina (Supplementary Fig. 3b).

**Figure 2.**
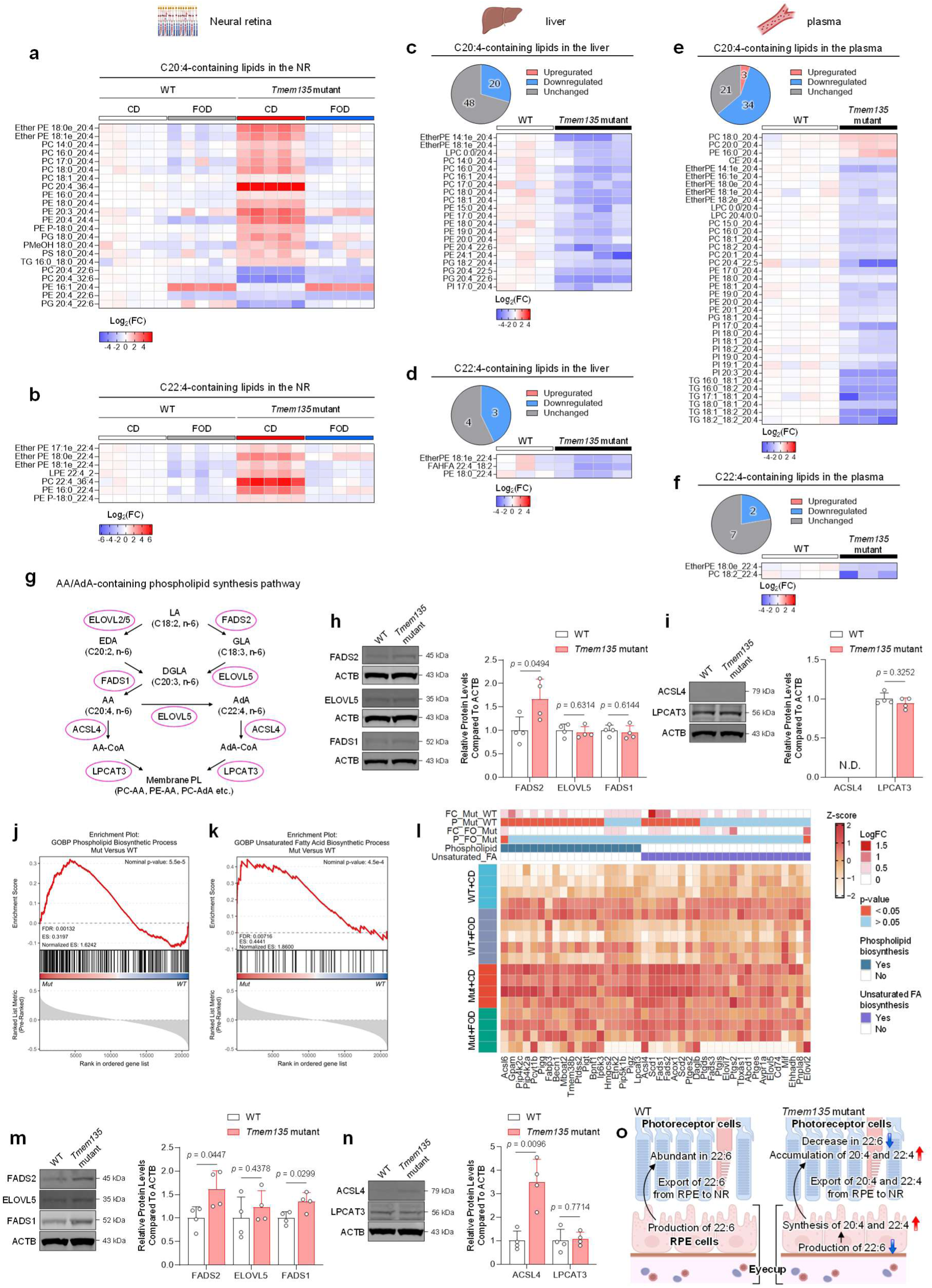
DHA deficiency drives AA/AdA phospholipid accumulation in the neural retina associated with an RPE biosynthetic response. **a**,**b**, Heatmaps of AA (20:4)-containing (**a**) and AdA (22:4)-containing (**b**) lipids in the neural retina of WT and *Tmem135* mutant mice fed a control diet (CD) or fish-oil diet (FOD). **c**,**d**, Pie charts and heatmaps of AA-containing (**c**) and AdA-containing (**d**) lipids in the liver of WT and *Tmem135* mutant mice. **e**,**f**, Pie charts and heatmaps of AA-containing (**e**) and AdA-containing (**f**) lipids in plasma from WT and *Tmem135* mutant mice. Pie charts in (**c–f**) show the numbers of lipid species that were increased, decreased, or unchanged in *Tmem135* mutant mice relative to WT mice. **g**, Schematic of the biosynthetic pathway for AA and AdA and their incorporation into phospholipids. **h**, Representative immunoblot and quantification of FADS2, ELOVL5 and FADS1 protein abundance in the neural retina. **i**, Representative immunoblot and quantification of ACSL4 and LPCAT3 protein abundance in the neural retina. **j**, GSEA enrichment plot for the phospholipid biosynthetic process gene set in *Tmem135* mutant versus WT eyecup. **k**, GSEA enrichment plot for the unsaturated fatty acid biosynthetic process gene set in *Tmem135* mutant versus WT eyecup. **l**, Heatmap showing expression of genes involved in the phospholipid and unsaturated fatty acid biosynthetic process. **m**, Representative immunoblot and quantification of FADS2, ELOVL5 and FADS1 protein abundance in the eyecup. **n**, Representative immunoblot and quantification of ACSL4 and LPCAT3 protein abundance in the eyecup. **o**, Schematic model of RPE AA/AdA biosynthesis and transfer to the neural retina during DHA deficiency. Data in (**h**,**i**,**m**,**n**) are presented as the mean ± s.d.; each data point represents an individual mouse (n = 4 per group). Immunoblot signals were normalized to ACTB. Sample sizes were n = 5 mice per group for neural-retinal lipidomics (**a**,**b**), n = 3–4 mice per group for liver and plasma lipidomics (**c–f**) and n = 4–5 mice per group for RNA-seq (**j–l**). In the pie charts (**c–f**), lipid species were classified as increased or decreased when *P* < 0.05 by a two-tailed unpaired Student’s *t*-test and as unchanged otherwise. Statistical significance in (**h**,**i**,**m**,**n**) was assessed using two-tailed unpaired Student’s *t*-tests. Exact *P* values are shown in the corresponding panels.

We next asked whether the accumulation of AA- and AdA-containing phospholipids in the neural retina reflected increased systemic availability of these fatty acids. In contrast to the neural retina and brain, AA- and AdA-containing phospholipids were predominantly reduced in the liver and plasma of *Tmem135* mutant mice (Fig. 2c–f). GC–FID-based fatty-acid profiling similarly showed increased AA and AdA in neural tissues but reduced levels in the liver and plasma (Supplementary Fig. 5a–d). Thus, long-chain ω-6 fatty acids accumulate in the neural retina despite their systemic decline, arguing against increased circulating availability as their source and supporting a locally regulated, neural tissue-selective compensatory response to DHA deficiency.

To investigate the basis of potential locally regulated neural tissue-selective ω-6 remodeling, we examined the pathway that converts linoleic acid (LA; 18:2) into AA and AdA and incorporates these fatty acids into phospholipids. This pathway involves FADS1 and FADS2, ELOVL2 and ELOVL5, and ACSL4, which activates AA to form AA-CoA for subsequent phospholipid incorporation (Fig. 2g). In the neural retina, ACSL4 abundance remained low in both WT and *Tmem135* mutant mice, and most enzymes involved in ω-6 PUFA synthesis and phospholipid incorporation showed no change (Fig. 2h,i). Thus, the accumulation of AA- and AdA-containing phospholipids could not be readily explained by increased biosynthesis within the neural retina itself. We therefore examined the adjacent RPE-containing eyecup as a potential source of the accumulating ω-6 fatty acids. Gene set enrichment analysis of eyecup RNA-seq data revealed coordinated enrichment of unsaturated fatty-acid biosynthesis and phospholipid metabolic pathways in *Tmem135* mutant mice (Fig. 2j–l). Consistent with this transcriptional signature, the eyecup exhibited abundant ACSL4 expression together with increased FADS1 and FADS2 protein abundance relative to WT controls (Fig. 2m,n).

Despite systemic DHA depletion, AA- and AdA-containing phospholipids did not increase uniformly across tissues. Instead, they were predominantly reduced in the liver and plasma but accumulated in the neural retina, indicating that retinal ω-6 enrichment was not driven by increased systemic availability. Moreover, the low abundance of ACSL4 and the absence of coordinated induction of enzymes required for long-chain ω-6 fatty-acid synthesis and phospholipid incorporation suggest that the neural retina itself has limited capacity to account for the increased production of AA- and AdA-containing phospholipids. Within the eye, retinal ω-6 accumulation coincided with an RPE-associated biosynthetic response and depletion of AA- and AdA-containing phospholipids in the eyecup. These opposing tissue-specific changes support a locally organized compensatory mechanism for retinal ω-6 lipid accumulation during DHA deficiency. The accompanying biosynthetic response in the RPE-containing eyecup raises the possibility that the RPE contributes ω-6 fatty acids to the neural retina (Fig. 2o).

### ω-6 lipid remodeling promotes oxidative lipid stress in the neural retina

The accumulation of AA- and AdA-containing phospholipids in the neural retina raised the possibility that compensatory ω-6 remodeling increases the burden of oxidized lipids during DHA deficiency. To test this possibility, we profiled oxidized derivatives of representative DHA-, AA-, and LA-containing phospholipids (1-palmitoyl-2-docosahexaenoyl-*sn*-glycero-3-phosphocholine (PDPC; PC 16:0_22:6), 1-palmitoyl-2-arachidonoyl-*sn*-glycero-3-phosphocholine (PAPC; PC 16:0_20:4), and 1-palmitoyl-2-linoleoyl-*sn*-glycero-3-phosphocholine (PLPC; PC 16:0_18:2)) were profiled in the neural retina. 1-palmitoyl-2-(4′-oxo-butanoyl)-*sn*-glycero-3-phosphocholine (POB-PC), oxidation products derived from PDPC, was markedly reduced, whereas oxidized species derived from ω-6 fatty acids, such as 1-palmitoyl-2-(5’-oxo-valeroyl)-*sn*-glycero-3-phosphocholine (POV-PC), 1-palmitoyl-2-succinoyl-*sn*-glycero-3-phosphocholine (PS-PC), 1-palmitoyl-2-glutaryl-*sn*-glycero-3-phosphocholine (PG-PC), and 1-palmitoyl-2-(9’-oxo-nonanoyl)-*sn*-glycero-3-phosphocholine (PON-PC), were broadly increased, with PAPC-derived oxidation products showing particularly robust elevation (Fig. 3a). Given the pronounced ω-6 remodeling toward AA and AdA observed in the neural retina, we next focused on abundant phosphatidylcholine species containing these fatty acids (PAPC, 1-stearoyl-2-arachidonoyl-*sn*-glycero-3-phosphocholine (SAPC; PC 18:0_20:4), 1-palmitoyl-2-adrenoyl-*sn*-glycero-3-phosphocholine (PAdPC; PC 16:0_22:4), and 1-stearoyl-2-adrenoyl-*sn*-glycero-3-phosphocholine (SAdPC; PC 18:0_22:4). Comprehensive profiling revealed a consistent increase in multiple oxidized AA-containing lipid species in the neural retina (Fig. 3b). In addition, 8-isoprostane—a non-enzymatic oxidation product derived from AA—was strongly increased and showed prominent accumulation in photoreceptor inner/outer segments (Fig. 3c). We next evaluated other oxidative stress in *Tmem135* mutant mice. Nuclear 8-hydroxy-2′-deoxyguanosine (8-OHdG), a major oxidative product of guanine, was strongly increased in photoreceptor nuclei of *Tmem135* mutant mice (Fig. 3d), indicating elevated oxidative stress. Markers of lipid peroxidation were likewise elevated. Immunostaining for acrolein-modified proteins revealed a diffuse increase throughout the neural retinas (Fig. 3e), and 4-hydroxynonenal (4-HNE) levels were markedly elevated in mutant mice (Fig. 3f). Importantly, the accumulation of 8-isoprostane and acrolein-modified proteins was substantially reduced by fish-oil supplementation, indicating that dietary restoration of DHA attenuates oxidative stress and lipid peroxidation in the mutant retina (Supplementary Fig. 6). In contrast, 4-HNE levels in the eyecup and the liver were unchanged (Supplementary Fig. 7a,b), consistent with the absence of AA accumulation in these tissues. SDHA, the mitochondrial complex II subunit containing a flavin site capable of ROS generation^26^, was also upregulated in mutant neural retinas (Supplementary Fig. 7c), whereas antioxidant enzymes CAT and GPX1 remained unchanged (Supplementary Fig. 7d).

**Figure 3.**
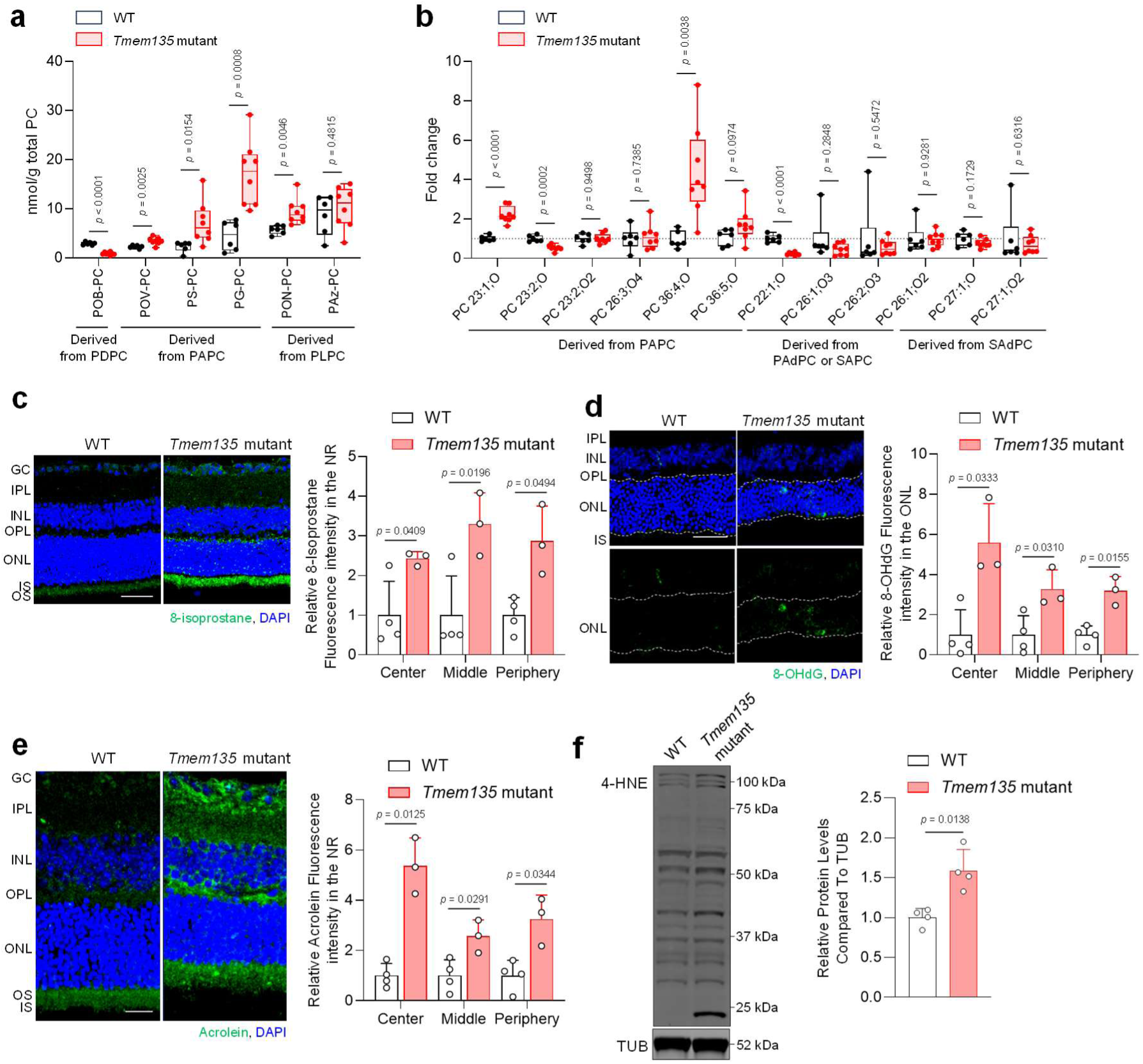
Compensatory ω-6 lipid remodeling promotes lipid peroxidation in the neural retina. **a**, Absolute abundance of oxidized lipid products derived from representative DHA-, AA- and LA-containing phosphatidylcholines in the neural retina. **b**, Relative abundance of oxidized lipid products derived from PAPC, SAPC, PAdPC and SAdPC in the neural retina. **c**, Representative retinal sections showing 8-isoprostane immunoreactivity, with quantification of relative fluorescence intensity in the neural retina. Scale bar, 50 μm. **d**, Representative retinal sections showing 8-OHdG immunoreactivity, with quantification of relative fluorescence intensity in the neural retina. Scale bar, 50 μm. **e**, Representative retinal sections showing acrolein-modified proteins, with quantification of relative fluorescence intensity in the neural retina. Scale bar, 25 μm. **f**, Representative immunoblot and quantification of 4-HNE-modified proteins in the neural retina. In (**a**,**b**), boxes show the median and interquartile range, whiskers indicate the minimum and maximum values, and individual data points represent individual mice. Data in (**c–f**) are presented as the mean ± s.d., with each data point representing an individual mouse. Sample sizes were n = 6–8 mice per group in (**a**,**b**), n = 3–4 in (**c–e**), and n = 4 in (**f**). Immunoblot signals in (**f**) were normalized to TUB. Statistical significance was assessed using two-tailed unpaired Student’s *t*-tests. Exact *P* values are shown in the corresponding panels.

Together, these findings demonstrate that compensatory ω-6 remodeling during DHA deficiency is accompanied by the preferential accumulation of oxidized ω-6 lipids, particularly AA-derived products, and a broader increase in oxidative damage within the neural retina. The reversal of these changes by DHA-rich fish-oil supplementation links the altered lipid environment to DHA availability.

### APOE-dependent oxidized-lipid clearance is coupled to Galectin-3 activation in subretinal microglia

Microglia are key cellular responders to oxidized lipids and can internalize these species for clearance^8^. The accumulation of oxidized ω-6 lipids in the neural retina and the migration of resident microglia into the subretinal space led us to investigate whether these microglia encounter oxidized phospholipids directly. Immunostaining with E06, an antibody that specifically recognizes oxidized phosphatidylcholine (oxPC), showed that in *Tmem135* mutant mice, E06 immunoreactivity was detected predominantly within subretinal microglia, with little detectable labeling in microglia residing in the IPL or OPL (Fig. 4a and Supplementary Fig. 8a). Moreover, E06 signals extensively colocalized with CD68, a lysosomal marker enriched in activated microglia (Fig. 4b), suggesting that oxidized phospholipids may accumulate within lysosomal compartments following their uptake by microglia. Oxidized lipid analysis revealed increased levels of representative oxidized ω-6–derived lipids in the eyecup with activated microglia (Fig. 4c and Supplementary Fig. 8b–g). Expanded profiling further revealed widespread accumulation of oxidized AA- and AdA-derived lipids in the eyecup, despite the depletion of their corresponding AA-containing phospholipid precursors, including PAPC and SAPC (Fig. 4d).

**Figure 4.**
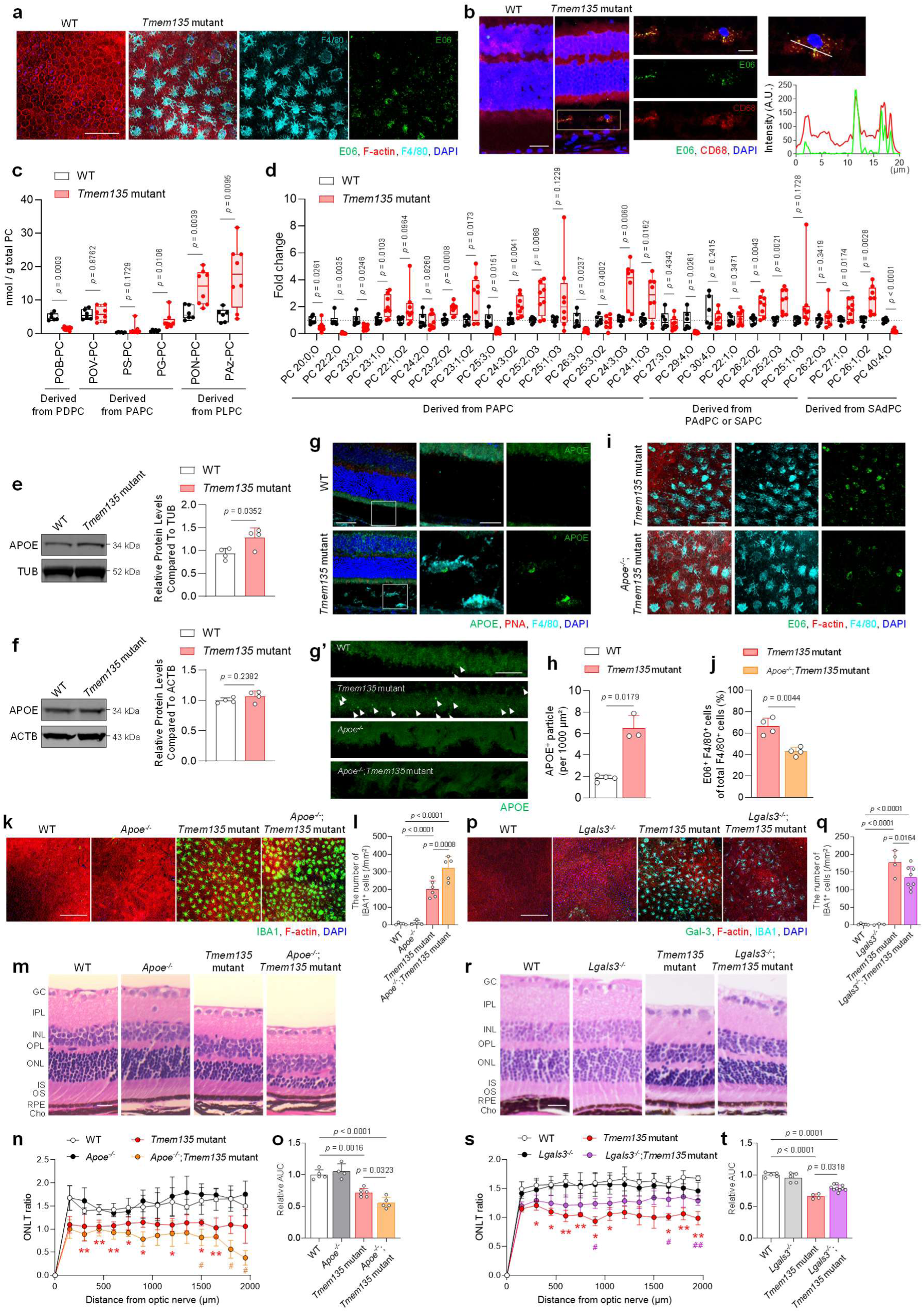
APOE-mediated oxidized-lipid clearance protects the retina, whereas Galectin-3 promotes progressive degeneration. **a**, Representative eyecup flat-mount images showing E06-reactive oxidized phospholipids (green) in IBA1⁺ subretinal microglia (cyan), with F-actin (red) and DAPI. Scale bar, 100 μm. **b**, Representative images showing colocalization of E06 immunoreactivity (green) with CD68 (red) in subretinal microglia. Scale bar, 25 μm. **c**, Absolute abundance of oxidized lipid products derived from representative DHA-, AA- and LA-containing phosphatidylcholines in the eyecup. **d**, Relative abundance of oxidized lipid products derived from PAPC, SAPC, PAdPC and SAdPC in the eyecup. **e**, Representative immunoblot and quantification of APOE protein abundance in the neural retina. **f**, Representative immunoblot and quantification of APOE protein abundance in the eyecup. **g**, Representative retinal sections showing APOE immunoreactivity (green) and F4/80⁺ subretinal microglia (cyan), with PNA labeling (red) of the photoreceptor inner/outer segment region and DAPI. Right, magnified view of the indicated region. Scale bars, 50 μm (left) and 15 μm (right). **g’**, Higher-magnification view of the photoreceptor inner/outer segment region. Scale bar, 20 μm. **h**, Quantification of APOE⁺ particles in the photoreceptor inner/outer segment region. **i**, Representative eyecup flat-mount images showing E06 immunoreactivity (green) in IBA1⁺ subretinal microglia (cyan) from *Tmem135* mutant and *Apoe*-deficient *Tmem135* mutant mice, with F-actin (red) and DAPI. Scale bar, 100 μm. **j**, Quantification of E06⁺ particles within subretinal microglia. **k**, Representative eyecup flat-mount images showing IBA1⁺ subretinal microglia (green) in WT, *Apoe*-deficient, *Tmem135* mutant and *Apoe*-deficient *Tmem135* mutant mice. Scale bar, 200 μm. **l**, Quantification of the number of IBA1⁺ subretinal microglia. **m**, Representative hematoxylin and eosin (H&E) staining of retinal sections. Scale bar, 25 μm. **n**, Outer nuclear layer thickness (ONLT) ratios. **o**, Area under the curve (AUC) of the ONLT ratio profiles. **p**, Representative eyecup flat-mount images showing IBA1⁺ subretinal microglia (green) in 6-month-old WT, *Lgals3*-deficient, *Tmem135* mutant and *Lgals3*-deficient *Tmem135* mutant mice. Scale bar, 200 μm. **q**, Quantification of the number of IBA1⁺ subretinal microglia. **r**, Representative H&E staining of retinal sections from 12-month-old WT, *Lgals3*-deficient, *Tmem135* mutant and *Lgals3*-deficient *Tmem135* mutant mice. Scale bar, 25 μm. **s**, ONLT ratios. **t**, AUC of the ONLT ratio profiles. In (**c**,**d**), boxes show the median and interquartile range, whiskers indicate the minimum and maximum values, and all individual data points are shown. Data in (**e**,**f**,**h**,**j**,**l**,**n**,**o**,**q**,**s**,**t**) are presented as the mean ± s.d. Each data point represents an individual mouse. Sample sizes were n = 6–8 mice per group in (**c**,**d**), n = 4 in (**e**,**f**), n = 3–4 in (**h**), n = 4 in (**j**), n = 4–6 in (**l**), n = 4–6 in (**n**,**o**), n = 3–8 in (**q**), and n = 4–10 in (**s**,**t**). Immunoblot signals in (**e,f**) were normalized to TUB or ACTB. Statistical significance was assessed using two-tailed unpaired Student’s *t*-tests (**c–f,h,j**), two-way ANOVA followed by Tukey’s multiple-comparisons test (**n,s**) or two-way ANOVA followed by Šídák’s multiple-comparisons test (**l,o,q,t**). Exact *P* values are shown in the corresponding panels. In (**n**), red asterisks indicate comparisons between WT and *Tmem135* mutant mice, whereas orange signs indicate comparisons between *Tmem135* mutant and *Apoe*-deficient *Tmem135* mutant mice. In (**s**), red asterisks indicate comparisons between WT and *Tmem135* mutant mice, whereas purple signs indicate comparisons between *Tmem135* mutant and *Lgals3*-deficient *Tmem135* mutant mice. */#*P* < 0.05, and**/##*P* < 0.01. Exact *P* values are shown in the remaining quantified panels; values below 0.0001 are reported as *P* < 0.0001.

Apolipoprotein E (APOE) regulates lipid transport and homeostasis in the central nervous system and its dysregulation contributes to age-related neurodegeneration^27^. APOE protein abundance was markedly increased in the neural retina of *Tmem135* mutant mice (Fig. 4e), whereas no significant difference was detected in bulk eyecup lysates (Fig. 4f). Within the eyecup, however, APOE immunoreactivity was spatially restricted to microglia located in the subretinal space. In the neural retina, APOE-containing particles accumulated within the photoreceptor inner and outer segments, and APOE-positive material was also detected within adjacent subretinal microglia (Fig. 4g,g′).

At the ultrastructural level, subretinal microglia contained abundant intracellular lipid-like deposits (Supplementary Fig. 8h). To test whether APOE mediates the transfer of oxidized lipids from the neural retina to subretinal microglia, we generated *Apoe*-deficient *Tmem135* mutant mice. *Apoe* deficiency markedly reduced E06-positive oxPC accumulation within subretinal microglia (Fig. 4i,j), while oxidized lipids accumulated further within the neural retina (Supplementary Fig. 9). This reciprocal distribution supports a model in which APOE-containing particles transport oxidized lipid cargo from the neural retina to subretinal microglia for uptake and clearance.

*Apoe* deficiency accelerated microglial recruitment to the subretinal space and exacerbated retinal degeneration in *Tmem135* mutant mice, despite the reduction in microglial oxPC accumulation (Fig. 4k–o). Together, these findings identify APOE as a mediator of oxidized-lipid transfer from the neural retina to subretinal microglia. Loss of this APOE-dependent transfer results in the retention and further accumulation of oxidized lipids within the neural retina, exacerbating retinal degeneration and increasing microglial recruitment to the subretinal space. These results support APOE-mediated trafficking of oxidized lipids to subretinal microglia as a protective retinal clearance pathway.

In parallel, we examined TREM2, a microglial receptor implicated in responses to tissue stress and lipid-associated signals^8,28^. *Trem2* transcript abundance was increased in both the neural retina and eyecup of *Tmem135* mutant mice, while TREM2 protein was increased in the eyecup and localized prominently to IBA1⁺ subretinal microglia (Supplementary Fig. 10a–c). Genetic deletion of *Trem2* reduced the number and size of subretinal microglia at 6 months but did not detectably improve retinal degeneration at this stage (Supplementary Fig. 10c–h). The decreased number of subretinal microglia by *Trem2* deficiency suggests that TREM2 contributes primarily to an early phase of microglial activation and subretinal accumulation. TREM2 may therefore mediate a parallel microglial response to stress signals arising from the neural retina. The lysosomal accumulation of oxidized phospholipids in subretinal microglia led us to examine galectin-3 (Gal-3 or LGALS3), a lysosomal stress-responsive lectin implicated in microglial inflammatory activation^29^. Gal-3 abundance was increased in the eyecup of *Tmem135* mutant mice and was highly expressed by IBA1⁺ microglia in the subretinal space (Supplementary Fig. 11a,b). Gal-3⁺ microglia also contained E06-positive puncta, linking Gal-3 induction to the intracellular accumulation of oxidized phospholipid-associated material (Supplementary Fig. 11c). Longitudinal analysis revealed that subretinal IBA1⁺ microglia accumulated between 2.5 and 6 months of age, remained elevated at 12 months and declined in the 15–18-month cohort (Supplementary Fig. 11d,e). Despite these changes in cell number, Gal-3 expression remained a persistent feature of subretinal microglia across all ages examined (Supplementary Fig. 11f). These cells also adopted progressively rounded morphologies and exhibited a modest reduction in cell size with age (Supplementary Fig. 11g,h), indicating sustained Gal-3-associated activation accompanied by progressive morphological remodeling.

To determine the functional contribution of Gal-3, we introduced *Lgals3* deficiency into the Tmem135 mutant background (Supplementary Fig. 12a,b). At 6 months, *Lgals3* deficiency modestly reduced subretinal microglial accumulation and cell size (Fig. 4p,q; Supplementary Fig. 12c,d) but produced only marginal attenuation of retinal degeneration (Supplementary Fig. 12e–g). At 12 months, the reduction in microglial cell size was no longer evident, although subretinal microglial accumulation remained modestly decreased (Supplementary Fig. 12h–j). Notably, retinal degeneration was more substantially ameliorated at this later stage (Fig. 4r–t). Together, these findings indicate that Gal-3 is not the primary driver of early microglial recruitment, but links sustained oxidized-lipid and lysosomal stress in subretinal microglia to the progressive phase of retinal degeneration.

Collectively, these findings delineate functionally distinct components of the subretinal microglial response to retinal stress: TREM2 contributes to early microglial activation and accumulation, APOE mediates the protective transfer and clearance of oxidized lipids, and sustained lipid-associated lysosomal stress engages Gal-3 to promote progressive retinal degeneration.

### DHA deficiency recapitulates age-related ω-6 lipid accumulation in the neural retina

It has been reported that DHA availability declines with aging and has been linked to age-related visual dysfunction^6^. Lipidomic profiling of aged mouse eyes revealed tissue-specific changes that closely mirrored those observed in *Tmem135* mutant mice. AA- and AdA-containing lipids increased broadly in the neural retina, while retinal DHA-containing phospholipids exhibited species- and class-dependent changes (Supplementary Fig. 13a–c). In the eyecup, most AA-, AdA- and DHA-containing lipids were concordantly reduced, although several PG species deviated from this overall pattern (Supplementary Fig. 13d–f). As PG is the immediate precursor of cardiolipin, these exceptions may reflect altered mitochondrial membrane remodeling during aging^30^. Although directionally similar, ω-6 remodeling encompassed a broader range of lipid species under DHA-deficient conditions than during physiological aging, suggesting that DHA deficiency amplifies this age-related response. Together, these findings indicate that DHA deficiency induces tissue-specific lipid remodeling that recapitulates and produces a more pronounced form of the aging-associated lipidomic state in both the neural retina and eyecup.

## Discussion

We identify a compensatory retinal lipid-remodeling response to DHA deficiency that increases ω-6 PUFA incorporation, oxidized-lipid burden, and retinal stress, driving age-related retinal phenotypes. In an attempt to alleviate this stress, the retina engages an APOE-dependent oxidized-lipid disposal pathway that facilitates the removal of oxidized lipids from the neural retina, while resident microglia migrate to the subretinal space and take up this lipid cargo. Sustained accumulation of oxidized lipids overwhelms the capacity of subretinal microglia to process them, leading to lysosomal stress and a galectin-3-dependent shift toward a degenerative microglial state that accelerates retinal degeneration. Thus, our findings support a model in which DHA deficiency initiates lipid-driven retinal stress and degeneration, whereas microglia initially engage in oxidized-lipid uptake but become maladaptive under sustained lipid burden, ultimately contributing to disease progression. The replacement of DHA-containing phospholipids by AA- and AdA-containing species also occurs during normal retinal aging, suggesting that the lipid-remodeling response identified here may represent a broader feature of the aging retina.

Notably, the resulting ω-6 enrichment in the neural retina did not reflect a systemic compensatory increase in ω-6 lipid availability, as AA- and AdA-containing phospholipids were predominantly reduced in the liver and plasma. Instead, the opposing changes in the RPE-containing eyecup and neural retina point to a locally organized compensatory response at the RPE–neural retina interface. Consistent with this model, DHA deficiency induced an RPE-associated transcriptional program involving PUFA biosynthesis and phospholipid remodeling, accompanied by increased FADS1, FADS2 and, particularly, ACSL4 protein abundance in the RPE-containing eyecup. By generating activated ω-6 fatty acids available for phospholipid incorporation, increased ACSL4 could contribute to the local production of AA- and AdA-containing lipids that subsequently accumulate in the neural retina. Thus, when DHA is limiting, the RPE may redirect local lipid supply toward alternative ω-6 PUFAs rather than relying on increased systemic ω-6 availability. The potential involvement of the RPE in neural retinal ω-6 lipid accumulation is consistent with its established role in photoreceptor lipid homeostasis. Photoreceptor outer segments undergo continuous renewal, yet a substantial fraction of their DHA is selectively recovered by the RPE after phagocytosis and recycled to photoreceptors, allowing the retina to conserve DHA despite this high membrane turnover^30,31^. The RPE therefore functions not merely as a conduit for circulating lipids but as an active metabolic hub that regulates PUFA availability to the outer retina. Our findings raise the possibility that this existing lipid-handling system also provides metabolic flexibility when DHA becomes limiting, redirecting local lipid metabolism and supply toward long-chain ω-6 PUFAs.

Compensatory AA and AdA incorporation helps maintain the highly polyunsaturated membrane composition required for photoreceptor function, but it may also increase susceptibility to lipid peroxidation. AA- and AdA-containing phospholipids are particularly prone to oxidation^7,33^, and their oxidation generates bioactive lipid products^34,35,36^. Consistent with this, DHA deficiency is linked to heightened retinal vulnerability to oxidative stress^18,19^, whereas endogenous DHA synthesis promotes photoreceptor survival under oxidative stress^37^. Thus, the accumulation of AA- and AdA-containing phospholipids during DHA deficiency provides an expanded pool of peroxidizable membrane substrates, offering a direct biochemical explanation for the preferential generation of ω-6-derived oxidized lipids in the mutant retina.

Our study suggests that oxidized lipids generated in the neural retina elicit a spatially organized microglial response. The retina provides a particularly tractable system for dissecting this process, because lipid-engaged microglia that migrate into the normally microglia-poor subretinal space can be distinguished from microglia that remain in the inner retina, allowing lipid sensing, migration and cargo handling to be resolved spatially. These subretinal microglia express lipid-sensing and lipid-handling markers that have also been reported in activated microglial states across other neurodegenerative conditions^27,38,39,40^, pointing to a shared lipid-responsive microglial program across neural tissues.

Our findings support a model in which DHA deficiency first generates a retinal stress state that precedes and drives the microglial response. DHA depletion was accompanied by oxidative stress, Müller glial activation and complement activation, all of which were attenuated by fish-oil supplementation. These changes indicate that disruption of retinal lipid homeostasis produces a broader tissue-stress response rather than acting initially through microglia themselves. Activated Müller glia associated closely with microglia in the neural retina and showed evidence of C3 involvement. Notably, *Edn2*, a photoreceptor-derived stress mediator that transmits stress signals to Müller glia^22^, was strongly upregulated and suppressed by fish-oil treatment, suggesting that photoreceptor stress is relayed to Müller glia through *Edn2*. These findings suggest a potential stress-responsive cascade in which photoreceptor stress activates Müller glia through *Edn2*, followed by complement signaling that promotes C3aR-dependent microglial activation and migration toward the outer retina and subretinal space. This interpretation is consistent with evidence linking Müller glia-derived C3 to subretinal immune-cell recruitment after retinal injury^23^ and C3aR signaling to microglial activation in response to oxidative retinal damage^24^. TREM2 appears to contribute to this early response, as *Trem2* deficiency reduced the subretinal microglial accumulation, suggesting that TREM2 helps microglia sense retinal stress-associated cues and mount an effective recruitment or retention response. Importantly, this occurred without detectable improvement in retinal degeneration at this stage, indicating that microglial accumulation itself is unlikely to be the primary cause of the early tissue damage. Instead, retinal stress directly compromises photoreceptors while simultaneously eliciting a microglial response that may help limit further damage, including through APOE-dependent transfer of oxidized lipids from the neural retina to subretinal microglia for clearance.

Once positioned in the subretinal space, microglia appear to participate in the clearance of oxidized lipids exported from the neural retina. Neural tissues can transfer oxidized lipids to glia through APOE-containing particles^41^, and loss of *Apoe* in our model impaired oxidized-lipid disposal and accelerated retinal dysfunction despite increased microglial accumulation. These findings support an early protective role for APOE-dependent lipid transfer and indicate that effective handling of lipid cargo, rather than microglial accumulation itself, limits retinal stress. TREM2, which recognizes oxidized lipids and APOE-associated ligands^8,42,43^, may further facilitate this lipid-handling response.

Persistent lipid uptake, however, appears to shift this lipid-handling response into a pathogenic state. Lysosomal dysfunction can drive disease-associated microglial states across neurodegenerative conditions^44^. Gal-3 is recruited to damaged lysosomes and coordinates lysosomal stress responses^45^, and has been linked to OxPC-associated neurodegeneration^46^ and microglia-mediated inflammation^29^. The late-stage protection afforded by *Lgals3* deletion therefore supports a model in which sustained oxidized-lipid loading progressively disrupts microglial lysosomal homeostasis, engaging a Gal-3-dependent program that accelerates retinal degeneration. This interpretation is consistent with studies in other neurodegenerative settings showing that microglial uptake of oxidized lipids can be protective^8^, although pathogenic effects have also been reported in other contexts^15,48^. Rather than indicating that oxidized-lipid-responsive microglia are uniformly protective or pathogenic, our findings suggest that their function changes with disease progression: early lipid uptake may limit retinal stress, whereas sustained lipid loading and lysosomal dysfunction progressively shift the same response toward a degenerative state. Thus, the consequences of microglial lipid handling may depend on both the duration and magnitude of lipid stress.

This adaptive-to-maladaptive transition from DHA to AA and AdA accumulation may be particularly relevant to retinal aging. Physiologically aged retinas showed PUFA remodeling in the same direction as that observed under DHA-deficient conditions, including increased AA- and AdA-containing lipids in the neural retina. Chronic DHA deficiency amplifies an age-related lipid-remodeling process sufficiently to expose its downstream consequences and thereby provides a means to dissect the sequence of events that may accompany impaired DHA homeostasis during aging. Viewed in this way, our data suggest that reduced DHA availability in the aging retina could increase reliance on compensatory AA/AdA incorporation, thereby promoting oxidized lipid burden and increasing the demand placed on APOE-dependent lipid disposal and microglial lysosomal processing. Pathology may emerge when this compensatory system can no longer match the chronic production of damaged lipids.

Our study has several limitations. Although our data support an RPE-associated source of compensatory AA/AdA synthesis, direct RPE-to-photoreceptor lipid flux remains to be demonstrated. In addition, the mechanism by which TREM2 promotes microglial migration into the subretinal space is unresolved, as is the fate of oxidized lipids when this microglial response is reduced.

Together, our findings define an adaptive-to-maladaptive lipid response to DHA deficiency. Compensatory AA/AdA remodeling increases oxidized-lipid production and initially engages protective APOE-dependent microglial disposal, but persistent lipid burden ultimately promotes a Gal-3-dependent degenerative response. Thus, retinal degeneration may emerge when compensatory lipid remodeling exceeds the capacity for safe lipid clearance and processing, providing a mechanistic framework for how disrupted PUFA homeostasis may contribute to retinal diseases characterized by lipid accumulation and chronic neuroinflammation.

## Methods

### Animal experiment

Generation of *Tmem135^FUN0^*^25^*^/FUN025^*mice (*Tmem135* mutants) on a C57BL/6J background was described previously^18^. The *Tmem135^FUN025/FUN025^*allele was maintained on a C57BL/6J background, and age- and sex-matched wild-type littermates or C57BL/6J mice (The Jackson Laboratory; [RRID: IMSR_JAX:000664]) were used as wild-type controls. *Apoe^-/-^* mice (B6.129P2-*Apoe^tm1Unc^*/J [RRID: IMSR_JAX:002052]), *Trem2^-/-^* mice (C57BL/6J-*Trem2^em2Adiuj^*/J [RRID: IMSR_JAX:027197]), and *Lgals3^-/-^* mice (B6.Cg-*Lgals3^tm1Poi^*/J [RRID: IMSR_JAX:006338]) were obtained from The Jackson Laboratory and crossed with *Tmem135^FUN025/FUN025^* mice to generate *Apoe^-/-^*;*Tmem135^FUN025/FUN025^* mice, *Trem2^-/-^*;*Tmem135^FUN025/FUN025^* mice, and *Lgals3^-/-^*;*Tmem135^FUN025/FUN025^* mice, respectively, on a C57BL/6J background. For microglial lineage tracing, *Rosa26^mTmG^* reporter mice (B6.129(Cg)-*Gt(ROSA)26Sor^tm4(ACTB-tdTomato,-EGFP)Luo^*/J [RRID: IMSR_JAX:007676]) were crossed with *Tmem119^CreERT2^*mice (C57BL/6-*Tmem119^em1(cre/ERT2)Gfng^*/J [RRID: IMSR_JAX:031820]) to generate *Tmem119^CreERT2/+^*;*Rosa^mTmG^*mice. These mice were subsequently crossed with *Tmem135* mutant mice to create *Tmem119^CreER/+^*;*Rosa^mTmG^*;*Tmem135^FUN025/FUN025^*mice. Male and female mice aged 2.5, 3, 6, 12 or 15–18 months were used as indicated for individual experiments. Mice were housed in the same animal facility at the University of Wisconsin–Madison under standardized conditions with a 12-h light–dark cycle (lights on at 06:00 and off at 18:00). Unless otherwise indicated, mice were maintained on a soy protein-free extruded rodent diet (#2020X; Inotiv). For fish-oil supplementation experiments, mice were fed a custom diet containing 10% (w/w) fish oil (#TD.230589; Inotiv) from weaning until tissue collection. To minimize circadian variation, tissue collection and associated experimental procedures were performed during the light phase within a consistent midday window (11:00–13:00). All animal procedures were conducted in accordance with the National Institutes of Health *Guide for the Care and Use of Laboratory Animals* and were approved by the Animal Care and Use Committee of the University of Wisconsin–Madison. Experimental design and reporting followed the ARRIVE guidelines.

### Sample collection and storage

For lipidomic analyses, mice were euthanized under isoflurane anesthesia; for all other experiments, mice were euthanized by CO₂ inhalation. Eyes were enucleated, and the cornea and lens were removed under a dissecting microscope to separate the neural retina from the eyecup. Following transcardial perfusion with PBS, the whole brain was collected. Tissues were flash-frozen in liquid nitrogen and stored at −80 °C until analysis.

### Sample preparation for lipidomic analysis

Neural retinas, eyecups and brains were stored at −80 °C until analysis and processed on dry ice. Tissue mass was measured to the nearest 0.01 mg using an analytical balance (XSR205; Mettler Toledo) and used for normalization. Tissues were transferred to PowerBead tubes (Qiagen; 13113-50) and extracted with 250 µl PBS, 225 µl methanol containing SPLASH LipidoMix internal standards (Avanti Polar Lipids; lot 3307-07; 10 µl per sample) and 750 µl methyl tert-butyl ether (MTBE). Samples were homogenized using a TissueLyzer II (Qiagen) at 30 Hz for three 30-s cycles, with 5 min on ice between cycles, followed by 15 min on ice. After centrifugation at 16,000 × *g* for 5 min at 4 °C, 500 µl of the upper organic phase was collected and dried in a SpeedVac concentrator (Savant). Lipid extracts were reconstituted in 150 µl isopropanol for LC–MS analysis. A process blank was prepared and analyzed alongside experimental samples. For each tissue, a pooled quality-control sample was generated by combining equal volumes of the reconstituted extracts and was used for lipid identification and signal normalization. Liver and plasma lipidomic data were obtained from a previously published dataset^19^ (Dryad, doi:10.5061/dryad.vx0k6djvm) and reanalyzed in the present study. Details of sample preparation and LC–MS acquisition for these samples have been described previously^19^.

### Untargeted LC-MS/MS-based lipidomic profiling

Lipid extracts were diluted in isopropanol at tissue- and ionization polarity-specific dilution factors before analysis. Lipids were separated on an Acquity UPLC BEH C18 column (1.7 µm, 2.1 × 100 mm; Waters) coupled to a VanGuard pre-column (1.7 µm, 2.1 × 5 mm; Waters) maintained at 50 °C, using an Agilent HiP 1290 Multisampler, 1290 Infinity II binary pump and column compartment coupled to an Agilent 6546 Accurate-Mass Q-TOF mass spectrometer equipped with a dual ESI source. For positive ion mode, the source gas temperature was 250 °C, drying gas flow was 12 L min^−1^, nebulizer pressure was 35 psig, VCap was 4,000 V, fragmentor voltage was 145 V, skimmer voltage was 45 V and octopole RF peak was 750 V. For negative ion mode, the corresponding settings were 350 °C, 12 L min^−1^, 25 psig, 5,000 V, 200 V, 45 V and 750 V. Reference masses (*m/z* 121.0509 and 922.0098 in positive mode; *m/z* 966.0007 and 112.9856 in negative mode) were continuously delivered to the second emitter of the dual ESI source at 15 µL min^−1^. Samples were analyzed in randomized order in separate positive- and negative-ionization runs over an *m/z* range of 100–1,500. Mobile phase A consisted of acetonitrile/water (60:40, v/v) containing 10 mM ammonium formate and 0.1% formic acid, and mobile phase B consisted of isopropanol/acetonitrile/water (90:9:1, v/v) containing the same additives. The gradient started at 15% B, increased to 30% B over 0–2.4 min, to 48% B over 2.4–3.0 min, to 82% B over 3.0–13.2 min and to 99% B over 13.2–13.8 min, followed by a hold at 99% B until 15.4 min and 4 min of re-equilibration at the initial conditions. The flow rate was 0.5 mL min^−1^ throughout. Injection volumes were 2 µL and 5 µL for positive- and negative-mode MS1 acquisition, respectively, and 4 µL and 7 µL for positive- and negative-mode MS/MS acquisition, respectively. MS/MS was performed using the same LC gradient with an isolation width of ∼1.3 *m/z* and a collision energy of 25 V. For each tissue, the pooled sample was analyzed in five iterative MS/MS runs with different precursor selections at each injection to increase coverage of lower-abundance lipid species.

### Lipid identification and quantification

Pooled MS/MS data and individual-sample MS1 data were processed using Agilent software. Lipids were identified from pooled MS/MS datasets using Lipid Annotator (Agilent) based on accurate precursor *m/z* values and experimental and theoretical fragment ions. Lipid class and acyl-chain composition were assigned when supported by the MS/MS data; otherwise, lipids were reported by summed composition, defined by total carbon number and degree of unsaturation. Separate identification libraries containing lipid species, *m/z* values and retention times were generated for positive- and negative-ionization modes. Lipids in individual samples were quantified from MS1 data using Profinder (Agilent). Ion chromatograms were extracted using the Lipid Annotator-derived libraries for protonated, deprotonated, ammonium-, sodium- or formate-adducted species, as appropriate for the ionization mode. Chromatographic peaks were integrated to determine lipid abundance, and the resulting data were exported for downstream statistical analysis.

### GC-FID-based fatty acid profiling

Total lipids were extracted from brain using the Folch method. Pentadecanoic acid (15:0) and heptadecanoic acid (17:0) were added to each sample as internal standards (25 nmol each). Lipid extracts were subjected to acid-catalyzed hydrolysis and methanolysis in 16% (v/v) concentrated HCl in methanol at 100 °C for 2 h to generate fatty acid methyl esters (FAMEs). FAMEs were extracted with hexane, purified by thin-layer chromatography and quantified using an Agilent 7890B gas chromatograph equipped with a flame ionization detector. Fatty acid composition was expressed as the mole percentage of each fatty acid species. Fatty acid profiling data for retina, plasma and liver were obtained from previously published raw data and reanalyzed for the present study^19^.

### LC–MS-based analysis of oxidized and PUFA-containing phospholipids

Neural retinas and eyecups were collected from 6-month-old mice. Frozen tissues were homogenized using a Macro Smash homogenizer in 1 ml of extraction solution (chloroform/methanol, 2:1, v/v) containing 100 µM butylated hydroxytoluene, 100 µM EDTA and SPLASH LIPIDOMIX Mass Spec Standard (1:500; Avanti Research). Homogenates were sonicated on ice for 5 min, incubated on ice for an additional 5 min and centrifuged at 6,000 × *g* for 10 min at 4 °C. The supernatant (600 µl) was mixed with 600 µl chloroform, 1000 µl methanol, and 1200 µl water and centrifuged at 2,000 × *g* for 20 min at 4 °C. The chloroform phase was collected, and the remaining sample was extracted twice more with 1200 µl chloroform. Combined chloroform fractions were dried under a stream of nitrogen gas, dissolved in methanol and stored at −80 °C until analysis. Lyso PC 18:1-d7 (1-oleoyl(d7)-2-hydroxy-*sn*-glycero-3-phosphocholine) or PC 15:0_18:1-d7 (1-pentadecanoyl-2-oleoyl(d7)-*sn*-glycero-3-phosphocholine) contained in SPLASH LIPIDOMIX was used as an internal standard. Lipid analysis was performed using an LCMS-8060 system equipped with an electrospray ionization source (Shimadzu), as previously described^49^. Chromatographic separation was performed on an InertSustain C18 column (2.1 × 150 mm, 3-µm particles; GL Sciences, Tokyo, Japan) maintained at 40 °C. The injection volume was 5 µl, and the autosampler was maintained at 4 °C. Mobile phase A consisted of 5 mM ammonium formate in acetonitrile/water (2:1, v/v), and mobile phase B consisted of isopropanol/methanol (19:1, v/v) containing 5 mM ammonium formate. The flow rate was 0.4 ml min^−1^. The proportion of mobile phase B was increased from 0 to 100% over 22.5 min, maintained at 100% until 27.5 min and then returned to 0% for 2.5 min of column re-equilibration under the initial conditions. Data were analyzed using Multi-ChromatoAnalysT v.1.3.3.0 (Beforce). Analyte abundances were calculated from peak-area ratios relative to the appropriate deuterated internal standard and normalized to total phospholipid content determined using the LabAssay™ Phospholipid kit (Catalog #295-94401; FUJIFILM Wako Pure Chemical Corporation). Authentic PS-PC and POB-PC standards (Avanti Research) and PG-PC, POV-PC, PON-PC and PAz-PC standards (Cayman Chemical) were used to support oxidized-PC peak assignments by comparison of retention times and chromatographic peak shapes. PAPC (Avanti Research), SAPC and SAdPC (Cayman Chemical) were oxidized as previously described^49^, and the resulting products were used as references for the retention times of oxidation-derived peaks. During the same analysis, non-oxidized phospholipids containing arachidonic acid (AA), adrenic acid (AdA) or docosahexaenoic acid (DHA) were measured concurrently by targeted LC–MS/MS. The monitored oxPC and non-oxidized phospholipid species, together with their corresponding MRM transitions and collision energies, are listed in Supplementary Tables 1 and 2, respectively. OxPC species for which no signal was detected in any sample were omitted from the figures.

### Histological analysis

Eyes were fixed in a solution containing 2% paraformaldehyde (PFA) and 2% glutaraldehyde overnight, followed by rinsing with phosphate-buffered saline (PBS) and embedding in paraffin. The paraffin-embedded samples were submitted to the Translational Research Initiatives in Pathology (TRIP) Core at the University of Wisconsin–Madison for tissue processing and sectioning. Serial sections (6 µm thick) were cut using a RM2135 microtome (Leica Microsystems, Wetzlar, Germany). Paraffin sections were stained with hematoxylin and eosin (H&E) according to standard protocols to visualize retinal layers, and images were acquired using an Axio Imager 2 microscope (Carl Zeiss MicroImaging, New York, USA) at 40× magnification.

### Electroretinograms

Full-field electroretinography (ERG) was performed in 6-month-old WT and *Tmem135* mutant mice fed either a control diet (CD) or a fish oil-supplemented diet (FOD) using an Espion E2 system (Diagnosys, Lowell, MA). The mice were dark-adapted overnight and anesthetized under dim red illumination by intraperitoneal injection of ketamine (80 mg/kg) and xylazine (16 mg/kg). The corneas were anesthetized with 0.5% proparacaine, the pupils were dilated with a drop of 1% tropicamide, and responses were recorded simultaneously from both eyes using corneal contact-lens electrodes. Hypromellose ophthalmic solution was applied to maintain corneal hydration and stable electrode contact. The reference and ground electrodes were placed subcutaneously in the cheek and tail, respectively, and the body temperature was maintained at 37 °C. Scotopic responses were recorded after dark adaptation. Photopic responses were recorded after 10 min of light adaptation to a 30 cd.s m^−2^ background. The flash intensities ranged from 0.03 to 30 cd.s m^−2^, with ten responses averaged at each intensity. The interstimulus intervals were 2 s for 0.03–3 cd s m^−2^ flashes and 5 s for 10–30 cd.s m^−2^ flashes. RPE function was assessed from c-wave responses elicited by 2.5 or 25 cd.s m^−2^ flashes using a 4-s acquisition period. Recordings were performed between 12:00 and 16:00, and all experimental groups were assessed within the same time window. ERG parameters were extracted using Espion V6 software.

### Western blot analysis

Neural retinas and eyecups were isolated separately, snap-frozen in liquid nitrogen and stored at −80 °C until use. Tissues were homogenized in RIPA buffer (#P189901, Thermo Fisher Scientific, Waltham, MA) supplemented with protease inhibitors (#11836170001, Thermo Fisher Scientific, Waltham, MA) using a Ultra-Turrax T8 tissue homogenizer. Protein concentrations were determined using a BCA Protein Assay Kit (#P123228, Thermo Fisher Scientific, Waltham, MA). Equal amounts of protein were aliquoted, reduced with XT Reducing Agent (#1610792, Bio-Rad, Hercules, CA) for 7 min at 105 °C, and resolved on 10% Bis-Tris Criterion XT gels (#3450112, Bio-Rad) in XT MOPS running buffer (#1610788, Bio-Rad) or XT MES running buffer (#1610789, Bio-Rad). Proteins were transferred onto Immun-Blot PVDF membranes (#1620177, Bio-Rad). Membranes were blocked with either 5% non-fat milk or EveryBlot Blocking Buffer (#12010020, Bio-Rad), followed by overnight incubation with primary antibodies at 4 °C. Each primary antibody was used at the dilution specified in Supplementary Table 3. After washing with TBST, membranes were incubated with the appropriate secondary antibodies (all diluted 1:5000): donkey anti-rabbit IgG 680RD (#926-68073, LI-COR), donkey anti-rabbit IgG 800CW (#926-32213, LI-COR), donkey anti-goat IgG 680RD (#926-68074, LI-COR), goat anti-mouse IgG1 800CW (#926-32350, LI-COR), and goat anti-mouse IgG2a 800CW (#926-32351, LI-COR). After final TBST washes, blots were visualized using the Odyssey Imaging System (LI-COR Biosciences, Lincoln, NE) and quantified with ImageJ software (NIH, Bethesda, MD). When required, membranes were stripped using NewBlot Stripping Buffer (LI-COR Biosciences) according to the manufacturer’s protocol and re-probed with additional primary antibodies. Band intensities were normalized to the corresponding loading controls.

### Immunohistochemistry of flat mount

Eyes were punctured at the cornea with a needle and fixed in 4% paraformaldehyde (PFA) for 30 min at room temperature. Samples were then immersed in 50% methanol for 15 min at room temperature, followed by 100% methanol at −20 °C for 15 min. After fixation, the cornea and lens were removed, and neural retinas were separated from the eyecups. Eyecups were blocked in 10% normal donkey serum for 30 min at room temperature, then incubated overnight at 4 °C with primary antibodies listed in Supplementary Table 3 under gentle agitation. After washing in PBS, samples were incubated for 2 h at room temperature with Alexa Fluor 488-conjugated (1:250; Thermo Fisher Scientific, Rockford, IL) and/or Cy3-conjugated (1:250; Jackson ImmunoResearch Laboratories, West Grove, PA) and/or Alexa Fluor 647-conjugated (1:250; Thermo Fisher Scientific, Rockford, IL) secondary antibodies. For F-actin staining, samples were incubated with Alexa Fluor 568–conjugated phalloidin (1:200; #A12380, Invitrogen, Carlsbad, CA) together with the secondary antibodies. Before mounting, four radial incisions were made to flatten the eyecup. Eyecup flat mounts were imaged using a Nikon A1R confocal microscope (Nikon Instruments, Tokyo, Japan) at 20× magnification. Microglial cell size was quantified using ImageJ by an investigator blinded to the experimental group.

### Immunohistochemistry of frozen-section

Eyes were fixed in 4% paraformaldehyde (PFA) for 2 h at 4 °C, cryoprotected in a graded sucrose series, and embedded in optimal cutting temperature (OCT) compound (Sakura Finetek USA, Torrance, CA). Cryostat sections (12 µm) were blocked in PBS containing 0.5% Triton X-100 and 2% normal donkey serum for 1 h at room temperature, then incubated overnight at 4 °C with primary antibodies listed in Supplementary Table 3. After washing in PBS, sections were incubated for 45 min at room temperature with Alexa Fluor 488-conjugated (1:250; Thermo Fisher Scientific, Rockford, IL) and/or Cy3-conjugated (1:250; Jackson ImmunoResearch Laboratories, West Grove, PA) and/or Alexa Fluor 647-conjugated (1:250; Thermo Fisher Scientific, Rockford, IL) secondary antibodies. Images were acquired using a Nikon A1R+ confocal microscope (Nikon Instruments, Melville, NY) equipped with GaAsP detectors, a high-speed resonant scanner. Image acquisition and analysis were performed using NIS-Elements AR software (Nikon Instruments).

### Bulk RNA-sequencing

Neural retinas and eyecups were collected separately from 6-month-old wild-type and *Tmem135* mutant mice of both sexes maintained on either the control or fish-oil diet. For each mouse, tissues from both eyes were pooled separately by tissue type. Samples were collected between 11:00 and 13:00, flash-frozen and submitted to GENEWIZ for RNA extraction, library preparation and sequencing. Total RNA was extracted using the RNeasy Plus Universal Mini Kit (Qiagen), quantified using a Qubit 2.0 Fluorometer (Life Technologies) and assessed for integrity using a 4200 TapeStation (Agilent Technologies). RNA-sequencing libraries were prepared using the NEBNext Ultra RNA Library Prep Kit for Illumina (New England Biolabs). Poly(A)+ RNA was enriched using oligo(dT) beads and fragmented at 94 °C for 15 min, followed by cDNA synthesis, end repair, 3′ adenylation, adapter ligation and limited-cycle PCR amplification. Libraries were assessed using an Agilent TapeStation and quantified by Qubit fluorometry and quantitative PCR. Libraries were sequenced using 150-bp paired-end reads on an Illumina. FASTQ files were generated and demultiplexed using bcl2fastq v.2.17. Raw sequencing data have been deposited in the Gene Expression Omnibus under accession number GSE348235.

### RNA sequencing data analysis

Raw gene-level read counts were analysed in R using DESeq2. Differential expression was modelled with experimental group as the design factor, and pairwise contrasts were performed as indicated. For gene set enrichment analysis (GSEA), genes were ranked by the DESeq2 Wald statistic, and preranked GSEA was performed using fgsea with gene sets obtained from MSigDB using msigdbr. Ensembl gene identifiers were mapped to mouse gene symbols using org.Mm.eg.db. For pathway-focused heatmaps, variance-stabilized expression values were standardized to gene-wise z scores.

### Microglial fate mapping

Microglial fate mapping was performed using *Tmem119^CreER/+^*;*Rosa^mTmG^*;*Tmem135^FUN025/FUN025^*mice. Tamoxifen-induced Cre recombination in *Tmem119*-expressing microglia switches the *Rosa26^mTmG^* reporter from membrane-targeted tdTomato to membrane-targeted GFP (mGFP), resulting in permanent labeling of these cells and their progeny. Beginning at 2 months of age, mice were fed tamoxifen-containing chow (#TD.130856; Inotiv) ad libitum for 1 week and then returned to the standard diet. Mice were analyzed at 3 or 6 months of age, corresponding to approximately 1- and 4-month chase periods, respectively. Eyecups were prepared as flat mounts, immunostained for IBA1 and imaged by confocal microscopy. Fate-mapped microglia were identified as mGFP^+^IBA1^+^ cells.

### Electron microscopy

Eyes were fixed with 2% paraformaldehyde (PFA) and 2% glutaraldehyde and submitted to the Electron Microscope Core at the University of Wisconsin-Madison for transmission EM processing as previously described. Eye sections were mounted on a 400-mesh thin bar grid, and images were collected where the grid bars intersected the neural retinas using a Phillips CM120 STEM microscope (FEI Company, Hillsboro, OR, USA) at 8,800X magnification. Mitochondria numbers were counted using NIH’s ImageJ software.

### Statistical analysis

RNA-sequencing analyses were performed in R as described above. All other analyses were performed using GraphPad Prism (GraphPad Software), except untargeted lipidomic analyses, which were conducted as described above. Two-group comparisons were performed using two-tailed unpaired Student’s *t*-tests. Multiple-group comparisons were performed using one-way ANOVA followed by Tukey’s test or two-way ANOVA followed by Tukey’s or Šídák’s test, as indicated in the figure legends. ERG data were analysed using two-way repeated-measures ANOVA with the Geisser–Greenhouse correction, followed by Šídák’s test of group marginal means. ONLT profiles were analysed using two-way ANOVA followed by Šídák’s test. Individual lipid species were compared using two-tailed unpaired Student’s *t*-tests and classified as altered at nominal *P* < 0.05 without correction for multiple testing. For microglial cell-size analyses, all cells within the acquired fields were quantified; mouse and cell numbers are reported in the figure legends. Data are presented as the mean ± s.d., except ERG data, which are presented as the mean ± s.e.m.; violin plots show the median and interquartile range. Exact *P* values are reported, with values below 0.0001 reported as *P* < 0.0001.

## ACKNOWLEDGMENTS

The authors thank the University of Wisconsin School of Medicine and Public Health Biomedical Research Model Services Shared Resource for use of its facilities and the Research Services team for their expertise in breeding the mice used in this study. The authors thank Toshi Kinoshita and the University of Wisconsin (UW) Translational Research Initiatives in Pathology laboratory (TRIP), supported by the UW Department of Pathology and Laboratory Medicine, UWCCC (P30 CA014520), and the Office of the Director, NIH (S10OD023526), for access to facilities and services, and Randall Massey and the University of Wisconsin Electron Microscope Core for tissue processing, sectioning and assistance with this study. We are grateful to the UW Biotechnology Center’s Advanced Lipidomics Platform for their time and effort in optimizing protocols for the lipidomics experiments. This work was supported by the Timothy William Trout Chair in Eye Research at the McPherson Eye Research Institute (to A. Ikeda); grants from the National Eye Institute (R01EY022086 to A. Ikeda and P30EY016665 to the Department of Ophthalmology and Visual Sciences at the University of Wisconsin–Madison, R01EY030513 to M.-P. Agbaga); JSPS KAKENHI (23H05481 and 25K24600 to K. Yamada); and grants from the Takeda Science Foundation (to K. Yamada), the Ono Medical Research Foundation (to K. Yamada), the Nagase Science and Technology Foundation (to K. Yamada) and the Hoansha Foundation (to K. Yamada); and an Unrestricted Grant from Research to Prevent Blindness, Inc. to the UW–Madison Department of Ophthalmology and Visual Sciences.

## AUTHOR CONTRIBUTIONS

Conceptualization – R.H., P.G., K.Y., and A.I.; Data curation – R.H., P.G., P.K.S., R.U., G.B.W., M.T., and R.S.B.; Formal analysis – R.H., P.G., P.K.S., R.U., G.B.W., M.L., M.T., R.S.B, J.W., and M.S.G.; Funding acquisition – M.P.A., K.Y., and A.I.; Investigation – R.H., P.G., P.K.S., R.U., and G.B.W.; Methodology – R.H., P.G., P.K.S., R.U., G.B.W., K.O., R.S.B., J.W., G.J.M., S.I., M.P.A., B.R.P., K.Y., and A.I.; Project administration – R.H., P.G., S.I., and A.I.; Resources – M.L., G.J.M., S.I., M.P.A., B.R.P., K.Y., and A.I.; Supervision – G.J.M., S.I., B.R.P., K.Y., and A.I.; Validation – R.H., P.G., P.K.S., R.U., G.B.W., K.O., S.I., M.P.A., B.R.P., K.Y., and A.I.; Visualization – R.H., P.G., and P.K.S.; Writing-original draft – R.H. and A.I.; Writing-review and editing – R.H., P.G., P.K.S., R.U., K.O., S.I., M.P.A., B.R.P., K.Y., and A.I.

