## Supplementary material for "DHA deficiency drives neural oxidized-lipid stress and microglial activation": Supplemenary figures

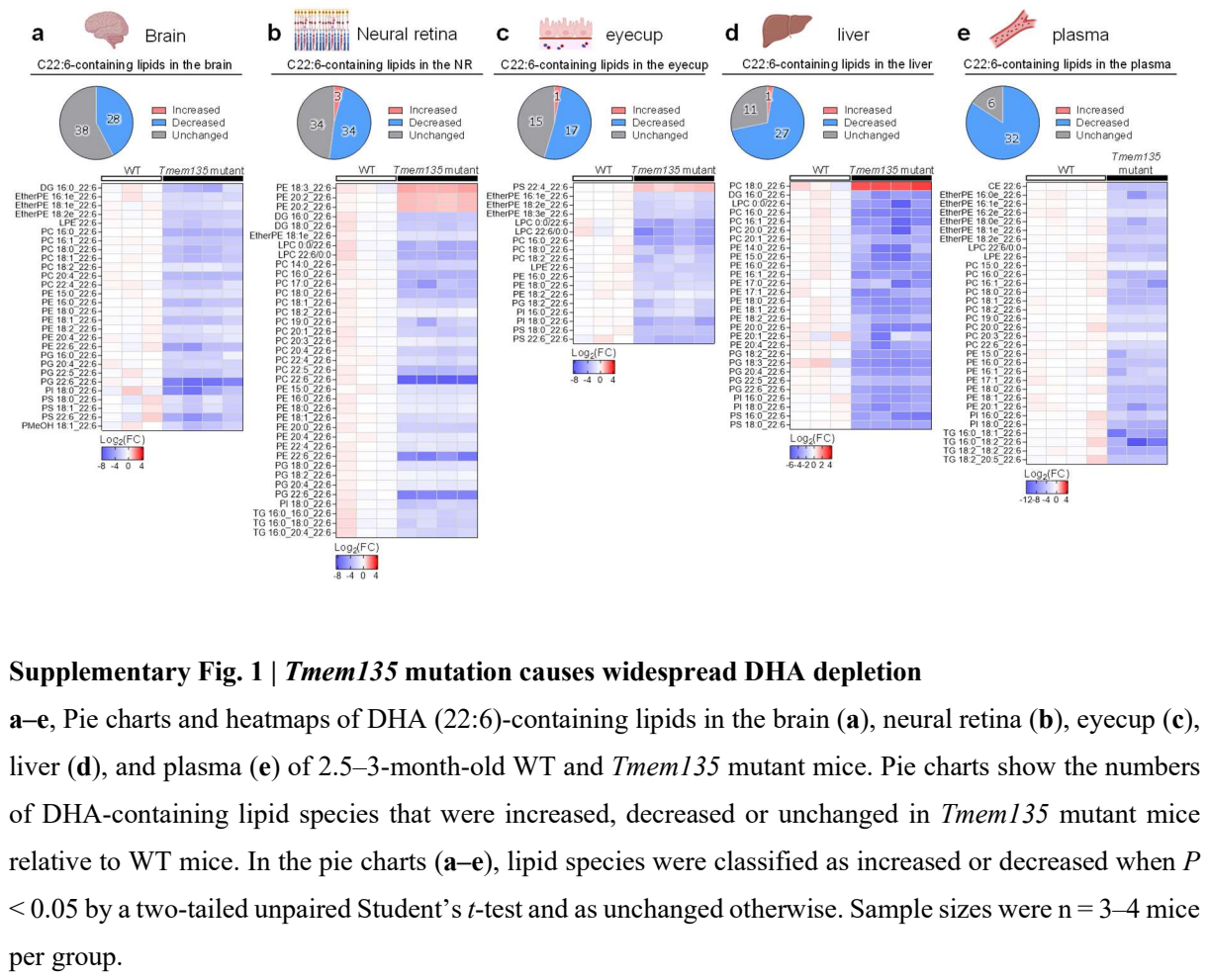

Supplementary Fig. 1 | Tmem135 mutation causes widespread DHA depletion

a–e, Pie charts and heatmaps of DHA (22:6)-containing lipids in the brain (a), neural retina (b), eyecup (c), liver (d), and plasma (e) of 2.5–3-month-old WT and Tmem135 mutant mice. Pie charts show the numbers of DHA-containing lipid species that were increased, decreased or unchanged in Tmem135 mutant mice relative to WT mice. In the pie charts (a–e), lipid species were classified as increased or decreased when P < 0.05 by a two-tailed unpaired Student’s t-test and as unchanged otherwise. Sample sizes were n = 3–4 mice per group.

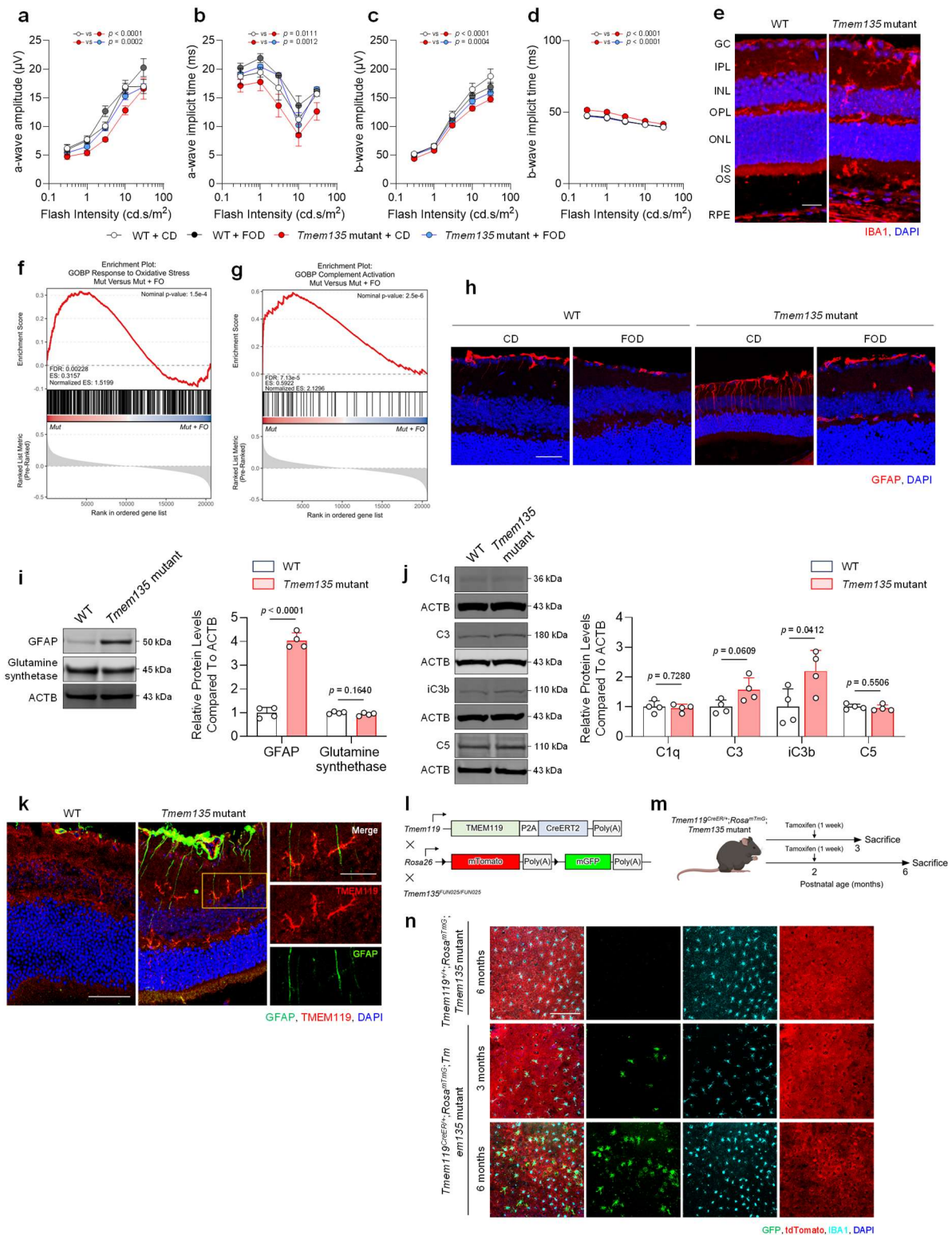

**Supplementary Fig. 2 | DHA deficiency engages a retinal stress–glial response and drives microglial migration**

**a–d**, Quantification of photopic ERG responses, showing a-wave amplitude (**a**), a-wave implicit time (**b**), b-wave amplitude (**c**), and b-wave implicit time (**d**). **e**, Representative retinal sections showing IBA1 immunoreactivity (red) and DAPI. Scale bar, 25 μm. **f**, GSEA enrichment plot for the response to oxidative stress gene set in neural retinas from *Tmem135* mutant mice fed a control diet versus a fish-oil diet. **g**, GSEA enrichment plot for the complement activation gene set in neural retinas from *Tmem135* mutant mice fed a control diet versus a fish-oil diet. **h**, Representative retinal sections showing GFAP immunoreactivity (red) and DAPI. Scale bar, 50 μm. **i**, Representative immunoblot and quantification of GFAP and glutamine synthetase protein abundance in the neural retina. **j**, Representative immunoblot and quantification of C1q, C3, iC3b, and C5 protein abundance in the neural retina. **k**, Representative retinal sections showing immunoreactivity for GFAP (green) and TMEM119 (red), with DAPI. Right, magnified view of the indicated region. Scale bars, 50 μm (left) and 25 μm (right). **l**, Schematic of the genetic strategy for fate mapping *Tmem119*-expressing microglia in *Tmem135* mutant mice using the *Tmem119*<sup>CreERT2</sup>;*Rosa26*<sup>mTmG</sup> reporter system. **m**, Timeline for tamoxifen induction and tissue collection in the microglial fate-mapping experiment. **n**, Representative eyecup flat mounts showing GFP-labelled IBA1<sup>+</sup> subretinal microglia (GFP, green; IBA1, cyan), with tdTomato (red) and DAPI. Scale bar, 200 μm. Data in (**a–d**) are presented as the mean ± s.e.m. For ERG analyses, responses from both eyes were averaged to obtain one value per mouse (n = 6–11 mice per group). Data in (**i,j**) are presented as the mean ± s.d.; each data point represents an individual mouse (n = 4 mice per group). Immunoblot signals in (**i,j**) were normalized to ACTB. For RNA-seq analyses (**f,g**), n = 4–5 mice per group, with neural retinas from both eyes of each mouse pooled. Statistical significance in (**a–d**) was assessed using two-way ANOVA with repeated measures on flash intensity and the Geisser–Greenhouse correction, followed by Šídák’s multiple-comparisons test of group marginal means. Statistical significance in (**i,j**) was assessed using two-tailed unpaired Student’s *t*-tests. Exact *P* values are shown in the corresponding panels; values below 0.0001 are reported as *P* < 0.0001.

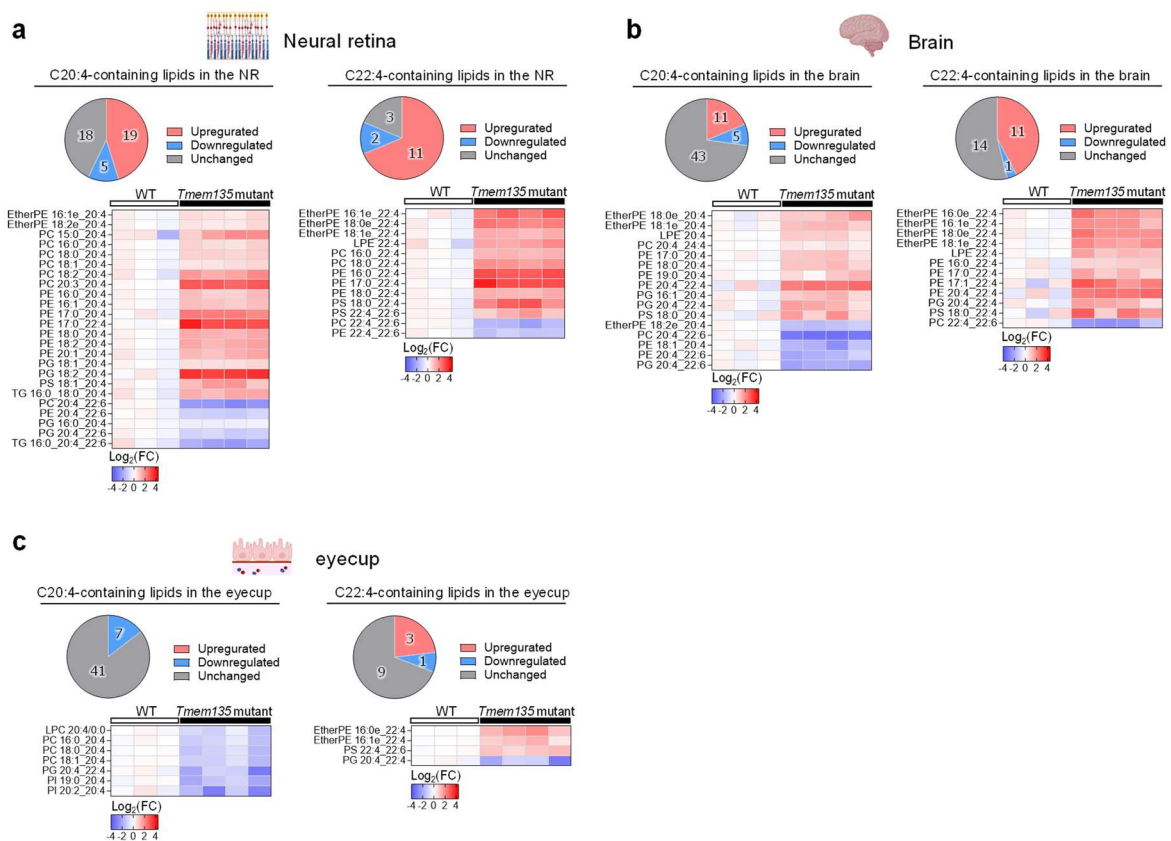

#### Supplementary Fig. 3 | DHA deficiency drives neural-selective accumulation of AA- and AdA-containing phospholipids

**a–c**, Pie charts and heatmaps of AA (20:4)- and AdA (22:4)-containing lipids in the neural retina (**a**), brain (**b**), and eyecup (**c**) of 2.5–3-month-old WT and *Tmem135* mutant mice. Pie charts show the numbers of AA- and AdA-containing lipid species that were increased, decreased or unchanged in *Tmem135* mutant mice relative to WT mice. In the pie charts (**a–c**), lipid species were classified as increased or decreased when $P < 0.05$ by a two-tailed unpaired Student's *t*-test and as unchanged otherwise. Sample sizes were $n = 3$–4 mice per group.

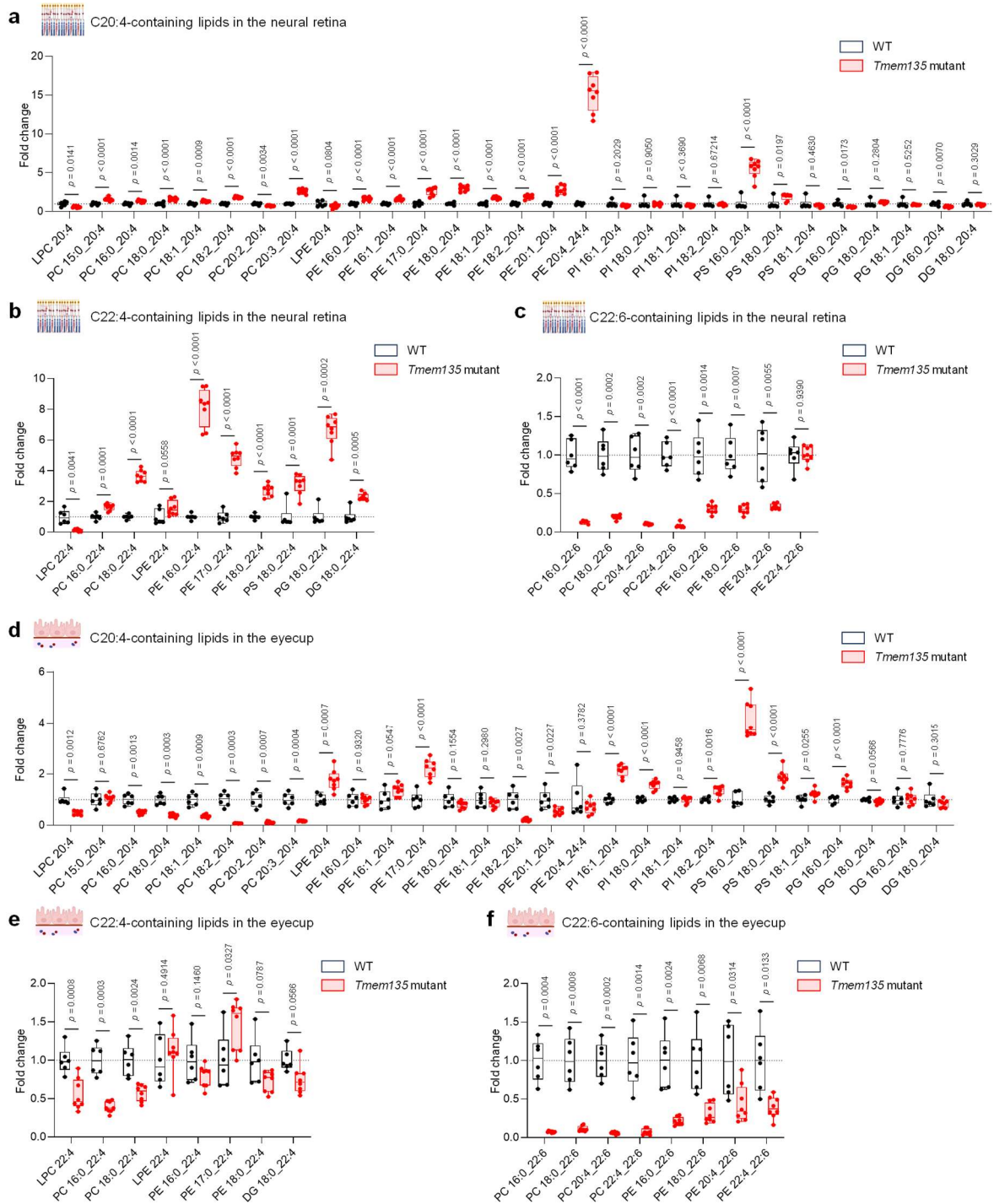

#### Supplementary Fig. 4 | DHA deficiency promotes AA/AdA accumulation in the neural retina with depletion in the eyecup

**a–c**, Quantification of AA (20:4)-containing (**a**), AdA (22:4)-containing (**b**) and DHA (22:6)-containing (**c**) phospholipids in the neural retina of 6-month-old mice. **d–f**, Quantification of AA (20:4)-containing (**d**), AdA (22:4)-containing (**e**) and DHA (22:6)-containing (**f**) phospholipids in the eyecup of 6-month-old mice. Boxes show the median and interquartile range, whiskers indicate the minimum and maximum values, and individual data points represent individual mice. $n = 6–8$ mice per group. Statistical significance was assessed using two-tailed unpaired Student's *t*-tests. Exact *P* values are shown; values below 0.0001 are reported as $P < 0.0001$.

**a**

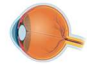

Neural retina & Eyecup

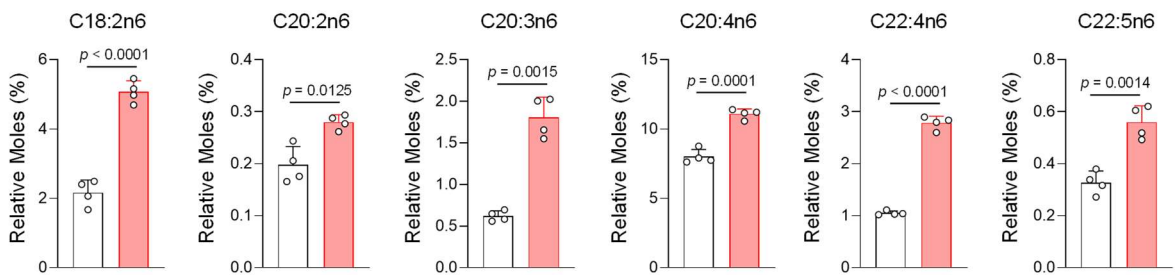

**b**

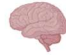

Brain

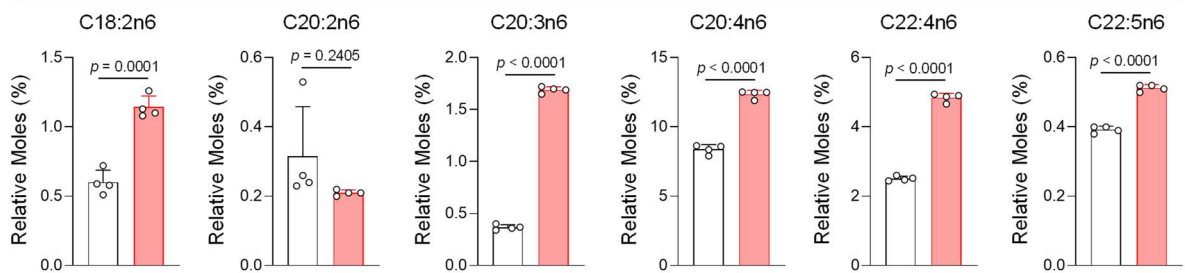

**c**

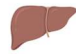

Liver

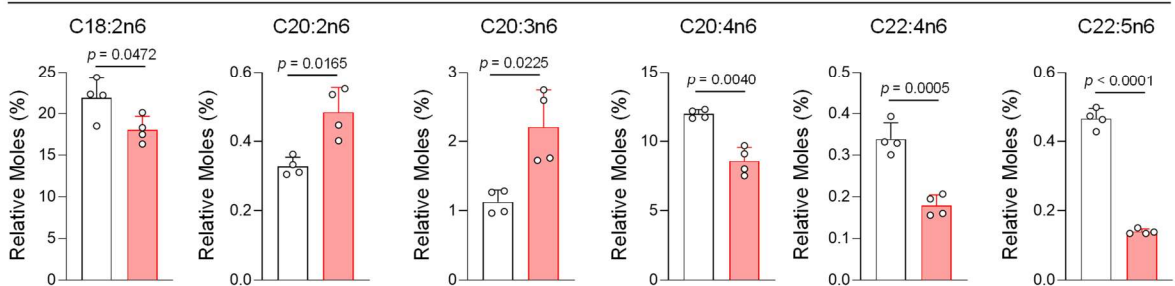

**d**

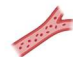

Plasma

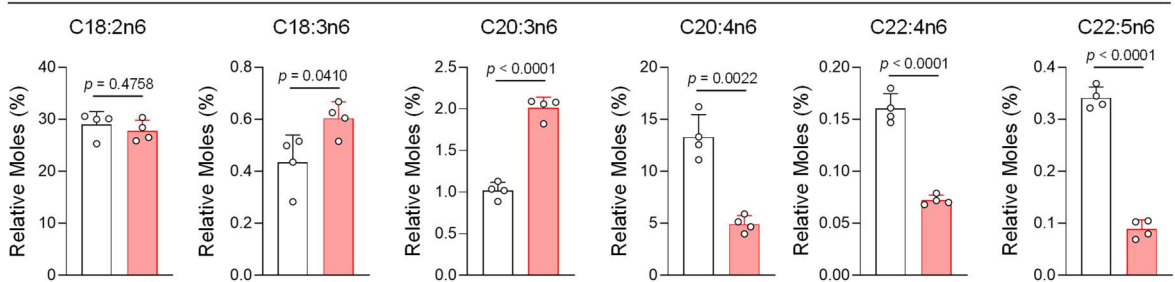

#### Supplementary Fig. 5 | DHA deficiency produces tissue-dependent changes in fatty-acid composition

**a–d**, GC–FID-based quantification of ω-6 fatty acids in the neural retina (**a**), brain (**b**), liver (**c**) and plasma (**d**) of WT and *Tmem135* mutant mice. Data are presented as the mean ± s.d.; each data point represents an individual mouse. $n = 4$ mice per group. Statistical significance was assessed using two-tailed unpaired Student's *t*-tests. Exact *P* values are shown; values below 0.0001 are reported as $P < 0.0001$.

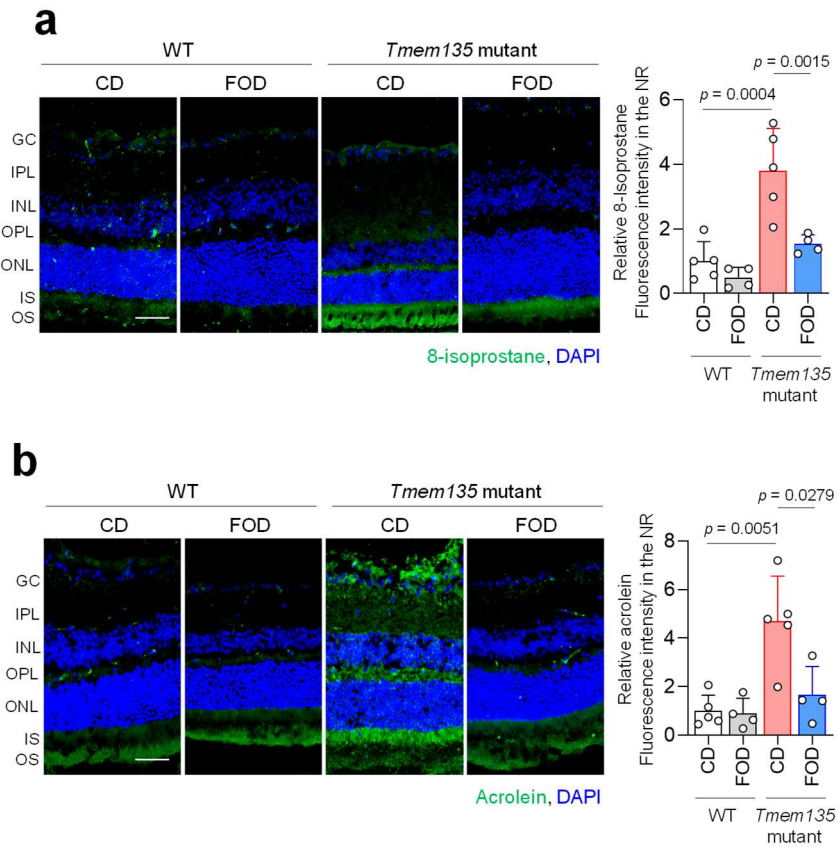

#### Supplementary Fig. 6 | DHA restoration suppresses lipid peroxidation in the neural retina

**a**, Representative retinal sections showing 8-isoprostane immunoreactivity, with quantification of relative fluorescence intensity in the neural retina. Scale bar, 50 $\mu\text{m}$. **b**, Representative retinal sections showing acrolein-modified proteins, with quantification of relative fluorescence intensity in the neural retina. Scale bar, 50 $\mu\text{m}$. Data are presented as the mean ± s.d.; each data point represents an individual mouse. $n = 4\text{--}5$ mice per group. Statistical significance was assessed using two-way ANOVA followed by Šidák's multiple-comparisons test. Exact $P$ values are shown in the corresponding panels.

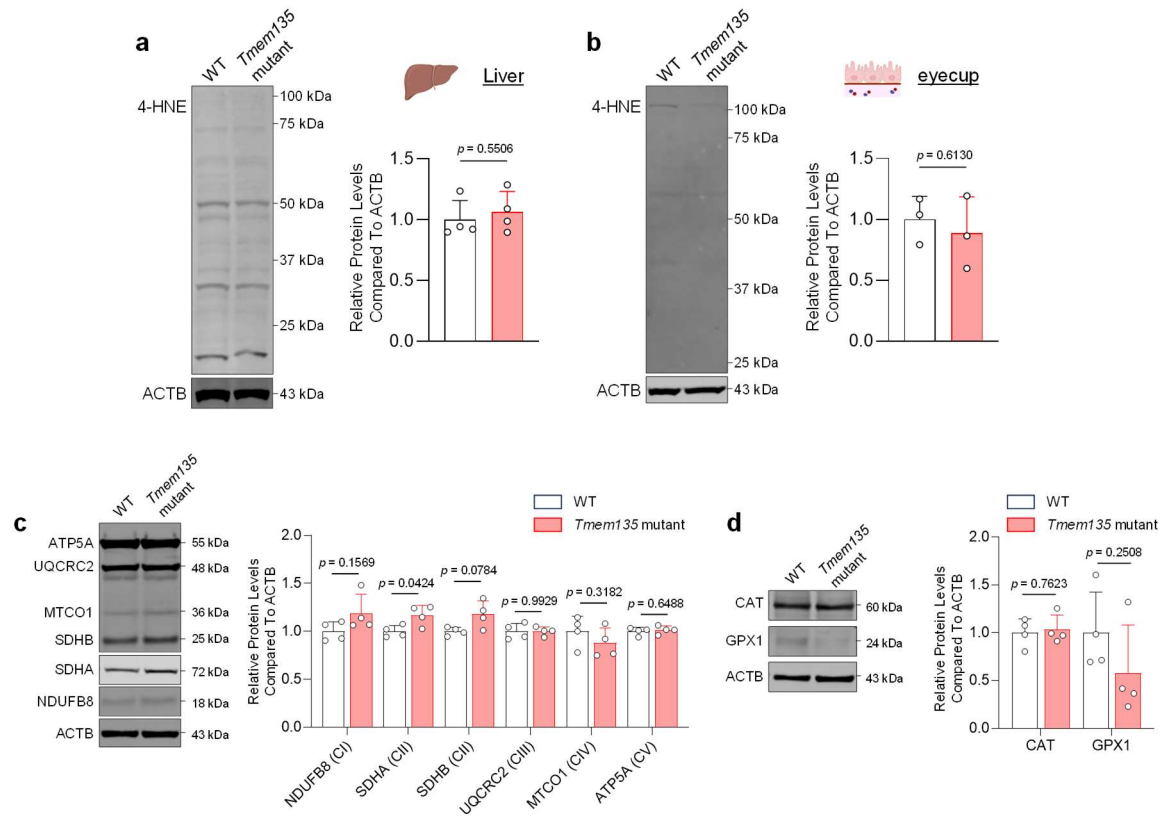

#### Supplementary Fig. 7 | Oxidative stress associated with DHA deficiency is selectively manifested in the neural retina

**a**, Representative immunoblot and quantification of 4-HNE-modified proteins in the eyecup. **b**, Representative immunoblot and quantification of 4-HNE-modified proteins in the liver. **c**, Representative immunoblots of NDUFB8 (complex I), SDHB and SDHA (complex II), UQCRC2 (complex III), MT-CO1 (complex IV) and ATP5A (complex V) in the neural retina, with quantification of protein abundance. **d**, Representative immunoblots of catalase (CAT) and glutathione peroxidase 1 (GPX1) in the neural retina with quantification of protein abundance. Protein abundance was normalized to ACTB. Data are presented as the mean ± s.d.; each data point represents an individual mouse. Sample sizes were $n = 4$ mice per group in **a,c,d** and $n = 3$ mice per group in **b**. Statistical significance was assessed using two-tailed unpaired Student's *t*-tests. Exact *P* values are shown.

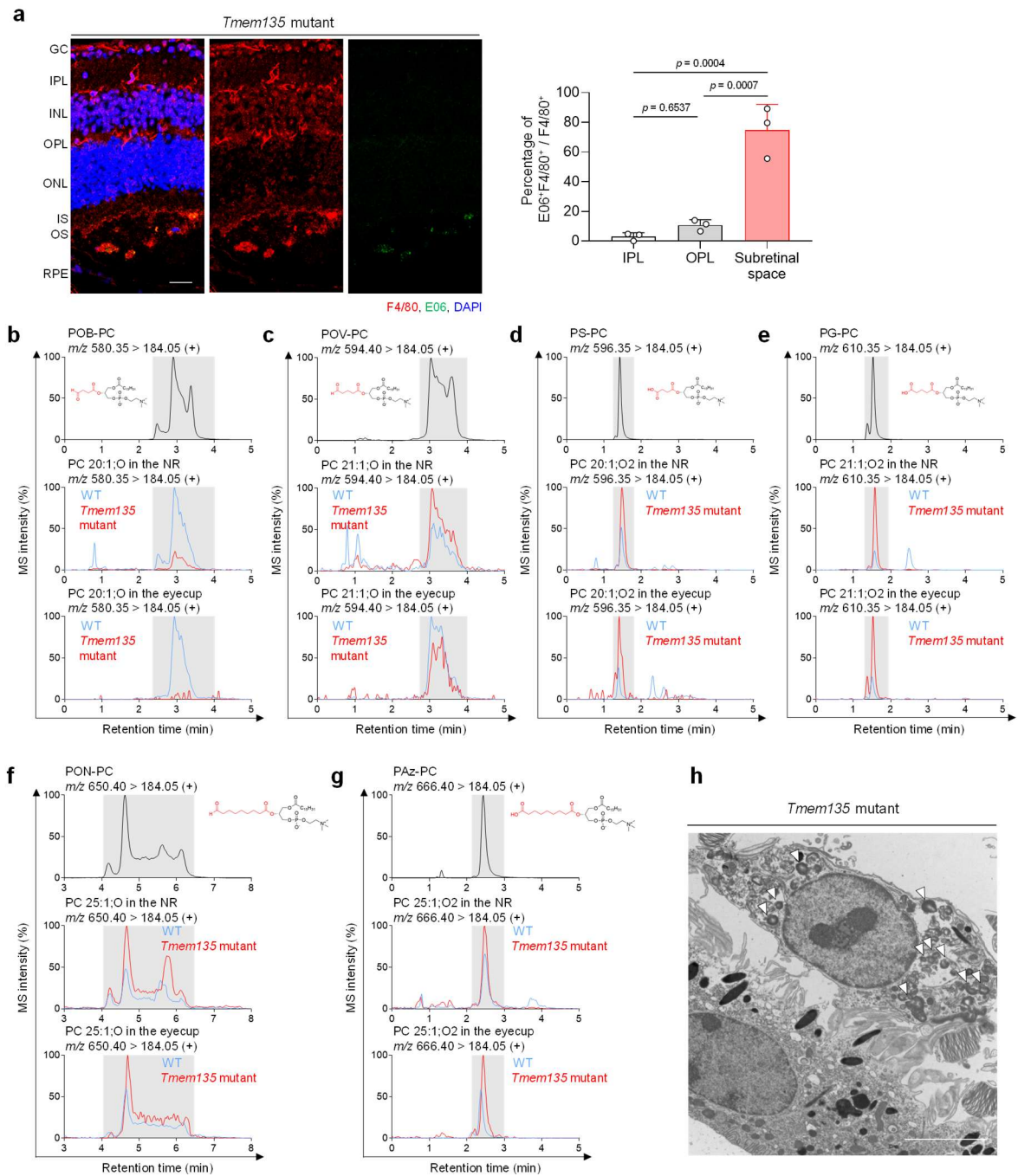

**Supplementary Fig. 8 | DHA deficiency drives oxidized-lipid generation in the neural retina and accumulation in the eyecup and subretinal microglia**

**a**, Representative retinal sections showing E06 immunoreactivity (green) and F4/80<sup>+</sup> microglia (red), with DAPI. Scale bar, 25 μm. **b–g**, Representative chromatograms of authentic standards and neural retina and eyecup samples from WT and *Tmem135* mutant mice for POB-PC (**b**), POV-PC (**c**), PS-PC (**d**), PG-PC (**e**), PON-PC (**f**) and PAz-PC (**g**). **h**, Representative electron micrograph of subretinal microglia. Arrowheads indicate intracellular lipid-like deposits. Scale bar, 5 μm. Data in (**a**) are presented as the mean ± s.d.; each data point represents an individual mouse (n = 3 mice). Statistical significance was assessed using one-way ANOVA followed by Tukey's multiple-comparisons test. Exact *P* values are shown.

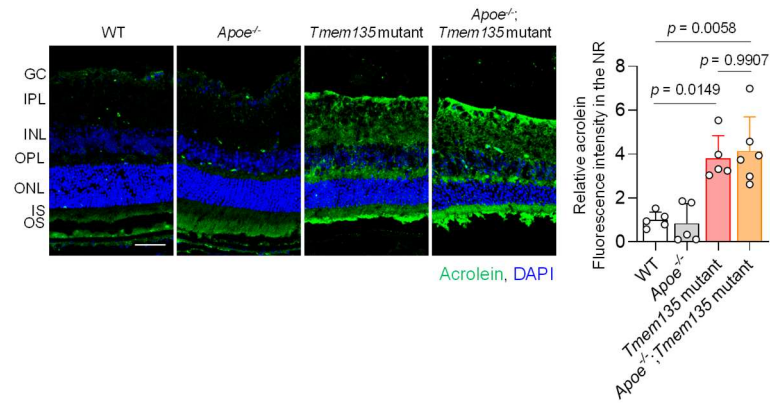

#### Supplementary Fig. 9 | Neural-retinal lipid peroxidation remains elevated in APOE-deficient *Tmem135* mutant mice

Representative retinal sections showing acrolein-modified proteins, with quantification of relative fluorescence intensity in the neural retina. Scale bar, 50 μm. Data are presented as the mean ± s.d.; each data point represents an individual mouse (n = 5–6 mice per group). Statistical significance was assessed using two-way ANOVA followed by Šidák's multiple-comparisons test. Exact *P* values are shown.

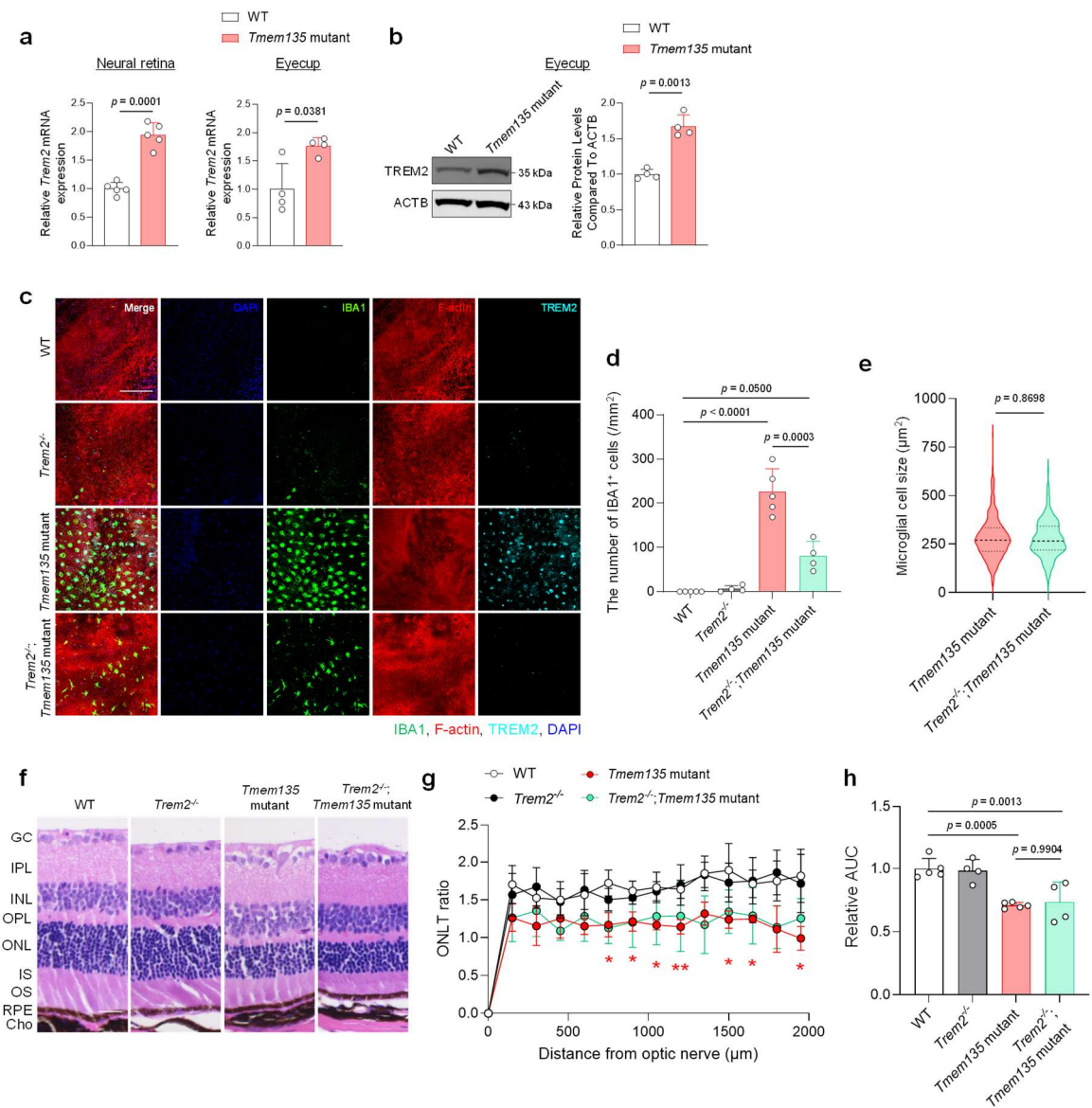

#### Supplementary Fig. 10 | TREM2 promotes subretinal microglial accumulation without affecting early retinal degeneration

**a**, Relative *Trem2* mRNA expression in the neural retina and eyecup. **b**, Representative immunoblot of TREM2 in eyecup lysates, with quantification of TREM2 protein abundance. **c**, Representative eyecup flat mounts showing IBA1<sup>+</sup> subretinal microglia from 6-month-old WT, *Trem2*-deficient, *Tmem135* mutant and *Trem2*-deficient *Tmem135* mutant mice. Scale bar, 200 μm. **d**, Quantification of the number of IBA1<sup>+</sup> subretinal microglia. **e**, Quantification of subretinal microglial cell size in *Tmem135* mutant and *Trem2*-deficient *Tmem135* mutant mice. **f**, Representative hematoxylin and eosin (H&E) staining of retinal sections from 6-month-old mice. Scale bar, 25 μm. **g**, Quantification of outer nuclear layer thickness (ONLT). **h**, Area under the curve (AUC) of ONLT profiles. Data in (a,b,d,g,h) are presented as the mean ± s.d.; each data point represents an individual mouse. Sample sizes were n = 4–5 mice per group in (a), n = 4 mice per group in (b) and n = 4–5 mice per group in (d,g,h). Protein abundance was normalized to ACTB. In (e), violin plots show the median and interquartile range; all IBA1<sup>+</sup> microglia within the acquired fields were quantified, comprising 444 and 197 cells from *Tmem135* mutant and *Trem2*-deficient *Tmem135* mutant mice, respectively, obtained from $n = 4\text{--}5$ mice per group. Statistical significance was assessed using two-tailed unpaired Student's *t*-tests (**a,b,e**), two-way ANOVA followed by Šídák's multiple-comparisons test (**d,h**) or two-way ANOVA followed by Tukey's multiple-comparisons test (**g**). In (**g**), red asterisks indicate comparisons between WT and *Tmem135* mutant mice; one and two asterisks denote $P < 0.05$ and $P < 0.01$, respectively. Exact *P* values are shown in the corresponding panels.

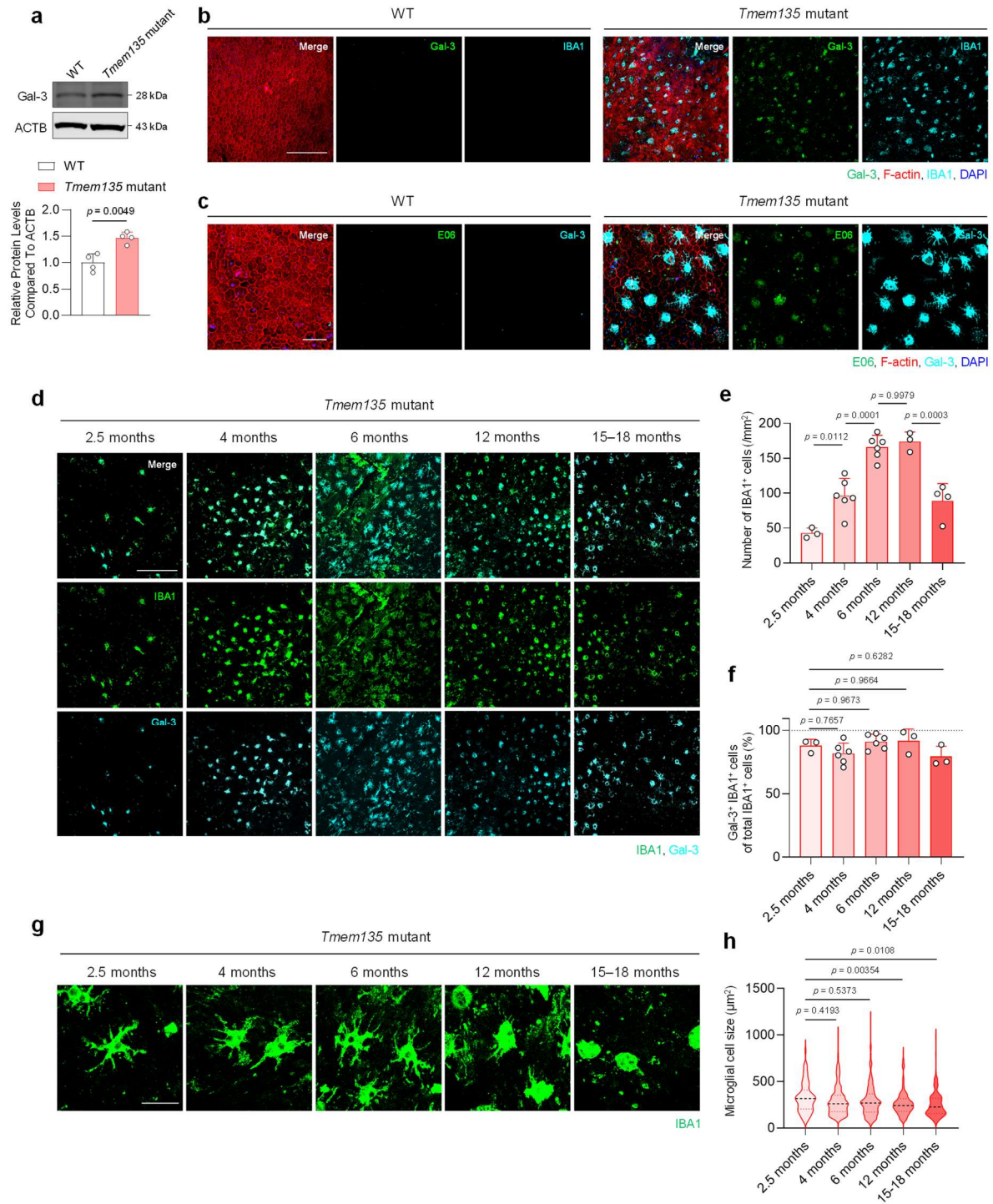

### Supplementary Fig. 11 | Galectin-3 marks a persistent subretinal microglial state across disease progression

**a**, Representative immunoblot of Galectin-3 (Gal-3) in eyecup lysates, with quantification of Gal-3 protein abundance. **b**, Representative eyecup flat mount showing Gal-3 immunoreactivity (green) in IBA1<sup>+</sup> subretinal microglia (cyan) with F-actin (red) and DAPI. Scale bar, 200 μm. **c**, Representative eyecup flat mount showing E06 immunoreactivity (green) and Gal-3 (cyan), with F-actin (red) and DAPI. Scale bar, 50 μm. **d**, Representative eyecup flat mounts showing Gal-3<sup>+</sup> IBA1<sup>+</sup> subretinal microglia in 2.5-, 4-, 6-, 12- and

15–18-month-old *Tmem135* mutant mice. Scale bar, 200 $\mu\text{m}$. **e**, Quantification of the number of IBA1<sup>+</sup> subretinal microglia. **f**, Quantification of the percentage of Gal-3<sup>+</sup> cells among IBA1<sup>+</sup> subretinal microglia. **g**, Representative higher-magnification images of subretinal microglia. Scale bar, 25 $\mu\text{m}$. **h**, Quantification of subretinal microglial cell size. Data in (**a,e,f**) are presented as the mean ± s.d.; each data point represents an individual mouse. Sample sizes were $n = 4$ mice per group in (**a**) and $n = 3\text{--}6$ mice per age group in (**e,f**). Protein abundance was normalized to ACTB. In (**h**), violin plots show the median and interquartile range; all IBA1<sup>+</sup> microglia within the acquired fields were quantified, comprising 74, 295, 400, 243, and 154 cells at 2.5, 4, 6, 12 and 15–18 months, respectively, obtained from $n = 3\text{--}6$ biologically independent mice per age group. Statistical significance was assessed using a two-tailed unpaired Student's *t*-test (**a**) or one-way ANOVA followed by Tukey's multiple-comparisons test (**e,f,h**). Exact *P* values are shown.

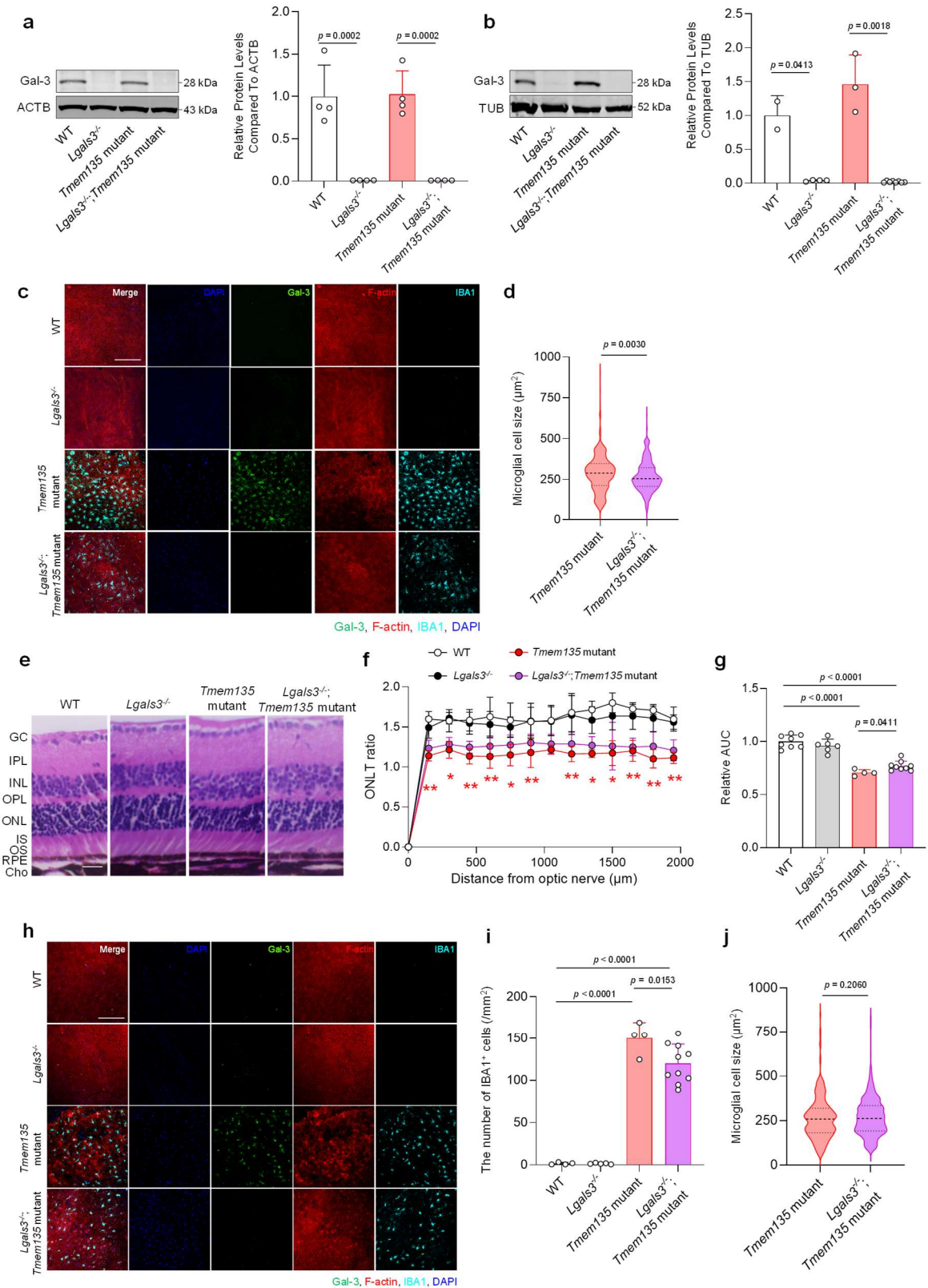

**Supplementary Fig. 12 | Galectin-3 deficiency attenuates subretinal microglial pathology and retinal degeneration**

**a**, Representative immunoblot of Galectin-3 (Gal-3) in neural retina lysates from WT, *Lgals3*-deficient,

*Tmem135* mutant and *Lgals3*-deficient *Tmem135* mutant mice, with quantification of Gal-3 protein abundance. **b**, Representative immunoblot of Gal-3 in eyecup lysates, with quantification of Gal-3 protein abundance. **c**, Representative eyecup flat mounts showing subretinal microglia in 6-month-old WT, *Lgals3*-deficient, *Tmem135* mutant and *Lgals3*-deficient *Tmem135* mutant mice. The same fields are shown in Fig. 4p, with the individual fluorescence channels presented separately. Scale bar, 200 μm. **d**, Quantification of subretinal microglial cell size at 6 months. **e**, Representative hematoxylin and eosin (H&E) staining of retinal sections from 6-month-old mice. Scale bar, 200 μm. **f**, Quantification of outer nuclear layer thickness (ONLT). **g**, Area under the curve (AUC) of ONLT profiles. **h**, Representative eyecup flat mounts showing subretinal microglia in 12-month-old WT, *Lgals3*-deficient, *Tmem135* mutant and *Lgals3*-deficient *Tmem135* mutant mice. Scale bar, 25 μm. **i**, Quantification of the number of subretinal microglia at 12 months. **j**, Quantification of subretinal microglial cell size at 12 months. Data in (**a**,**b**,**f**,**g**,**i**) are presented as the mean ± s.d.; each data point represents an individual mouse. Sample sizes were $n = 4$ mice per group in (**a**), $n = 2-8$ mice per group in (**b**), $n = 4-9$ mice per group in (**f**,**g**) and $n = 4-10$ mice per group in (**i**). In (**d**,**j**), violin plots show the median and interquartile range; all IBA1<sup>+</sup> microglia within the acquired fields were quantified. In (**d**), 345 and 410 cells were analysed from *Tmem135* mutant and *Lgals3*-deficient *Tmem135* mutant mice, respectively; in (**j**), 261 and 475 cells were analysed, respectively. Cells were obtained from $n = 4-10$ mice per group. Statistical significance was assessed using two-tailed unpaired Student's *t*-tests (**d**,**j**) or two-way ANOVA followed by Šidák's multiple-comparisons test (**a**,**b**,**f**,**g**,**i**). In (**f**), red asterisks indicate comparisons between WT and *Tmem135* mutant mice; one and two asterisks denote $P < 0.05$ and $P < 0.01$, respectively. Exact *P* values are shown in the corresponding panels.

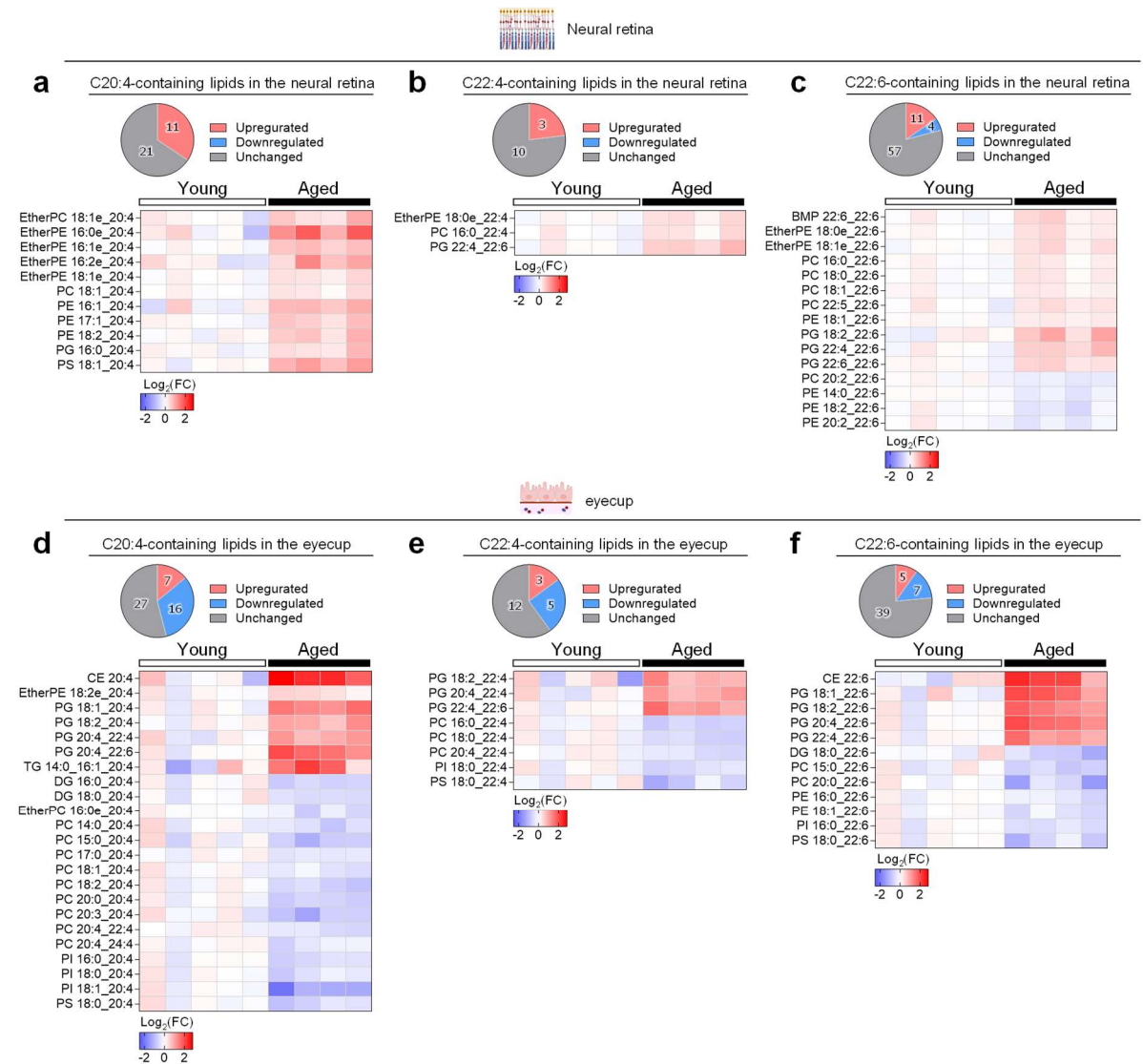

**Supplementary Fig. 13 | Physiological aging is associated with ω-6 lipid accumulation in the neural retina and PUFA depletion in the eyecup**

**a–c**, Pie charts and heatmaps of AA (20:4)-containing (**a**), AdA (22:4)-containing (**b**) and DHA (22:6)-containing (**c**) lipids in the neural retina of 2-month-old (young) and 17-month-old (aged) WT mice. **d–f**, Pie charts and heatmaps of AA (20:4)-containing (**d**), AdA (22:4)-containing (**e**) and DHA (22:6)-containing (**f**) lipids in the eyecup of 2-month-old (young) and 17-month-old (aged) WT mice. In the pie charts (**a–f**), lipid species were classified as increased or decreased when $P < 0.05$ by a two-tailed unpaired Student's $t$-test and as unchanged otherwise. Sample sizes were $n = 4–5$ mice per age group.
